# Transiently structured head domains control intermediate filament assembly

**DOI:** 10.1101/2020.03.27.012112

**Authors:** Xiaoming Zhou, Yi Lin, Glen Liszczak, Vasily Sysoev, Lillian Sutherland, Eiichiro Mori, Dylan Murray, Masato Kato, Robert Tycko, Steven L. McKnight

**Affiliations:** Department of Biochemistry, UT Southwestern Medical Center, 5323 Harry Hines Blvd. Dallas, TX 75390; Department of Future Basic Medicine, Nara Medical University, 840 Shijo-cho, Kashihara Nara, Japan; Department of Chemistry, University of California, Davis, California 95616; Department of Chemical Physics, NIDDK, National Institutes of Health, Bethesda, Maryland 20814

**Keywords:** low complexity domains, intermediate filaments, intrinsically disordered proteins, phase separation, human genetic mutations, labile cross-β structures, dynamic cellular assemblies, solid state NMR, segmental isotope labeling, *in situ* structural analysis

## Abstract

Low complexity (LC) head domains 92 and 108 residues in length are, respectfully, required for assembly of neurofilament light (NFL) and desmin intermediate filaments (IFs). As studied in isolation, these IF head domains interconvert between states of conformational disorder and labile, β-strand-enriched polymers. Solid state nuclear magnetic resonance (ss-NMR) spectroscopic studies of NFL and desmin head domain polymers reveal spectral patterns consistent with structural order. A combination of intein chemistry and segmental isotope labeling allowed preparation of fully assembled NFL and desmin IFs that could also be studied by ss-NMR. Assembled IFs revealed spectra overlapping with those observed for β-strand-enriched polymers formed from the isolated NFL and desmin head domains. Phosphorylation and disease causing mutations reciprocally alter NFL and desmin head domain self-association, yet commonly impede IF assembly. These observations show how facultative structural assembly of LC domains via labile, β-strand-enriched self-interactions may broadly influence cell morphology.

## Introduction

Mammalian genomes encode 70-80 unique proteins that assemble into a variety of intermediate filaments. These filaments play cyto-architectural roles that vary considerably in skin, muscle and nerve cells (Alberts, 2002). Fluorescence recovery after photo-bleaching (FRAP) experiments have shown that intermediate filaments can be surprisingly dynamic, allowing insertion of new polypeptide subunits internal to the longitudinal axis of existing filaments (Vikstrom et al., 1992). Despite access to a wealth of information pertinent to the form and function of intermediate filaments, the processes controlling their assembly, disassembly and regulation remain mysterious.

Intermediate filaments are defined by centrally located α-helical segments 300-350 residues in length. These central, α-helical segments are flanked on either end by head and tail domains thought to be devoid of structural order (Herrmann and Aebi, 2016; Kornreich et al., 2015). Assembly of IFs begins with the parallel association of two α-helical segments to form a coiled-coil. Pairs of coiled-coil dimers, in turn, associate in an anti-parallel orientation to form tetramers. Dimer and tetramer formation of IF proteins can proceed in the absence of their intrinsically disordered head and tail domains. By contrast, the end-to-end joining of tetramers into long proto-filaments, as well as the assembly of eight protofilaments around one another to form mature, 10 nm IFs, cannot proceed with variants trimmed of head and tail domains.

The intrinsically disordered head and tail domains of IF proteins are of low sequence complexity (Figure S1). Studies of the isolated head domains of vimentin, peripherin, internexin, and the light, medium, and heavy neurofilament isoforms have given evidence of phase separation in the form of gel-like condensates (Lin et al., 2016). In all six cases, head domains were observed to become phase separated upon incubation at high concentration. By contrast, similar treatment of the isolated tail domains of the same six proteins failed to affect phase separation. Longstanding studies of many different IF proteins have shown that maturation of 10 nm filaments is dependent upon an intrinsically disordered head domain (Ching and Liem, 1999; Gill et al., 1990; Petzold, 2005). It has been postulated that the head domain interactions leading to phase separation might be instructive as to how these intrinsically disordered protein domains assist in the assembly of intermediate filaments (Lin et al., 2016). Here we describe experiments designed to investigate this hypothesis.

## Results

The heavy, medium, and light neurofilament isoforms are important for the morphological integrity of axons, where they interact intimately with longitudinally oriented microtubules and exist at an abundance ten-fold greater than microtubules or actin filaments (Yuan et al., 2017). The studies reported herein were initially focused on the neurofilament light (NFL) isoform. The head domain of the NFL isoform, 92 residues in length, was prepared as a green fluorescent protein (GFP) fusion protein containing an amino-terminal 6xHis-tag, expressed in bacterial cells, purified, and incubated at high concentration under physiologic conditions of salt and pH (Experimental Procedures). As described previously, these procedures led to the formation of a gel-like condensate composed of uniform, amyloid-like polymers (Lin et al., 2016).

GFP:NFL head domain hydrogel droplets 5 mm in diameter were formed on parafilm and incubated for 2hr at 4°C with a soluble extract prepared from mouse brain tissue (Experimental Procedures). Following binding, hydrogel droplets were recovered by centrifugation, washed twice with phosphate-buffered saline, and melted by exposure to binding buffer supplemented with 6 M guanidine-HCl. Soluble material was passed through a nickel affinity resin to remove the 6xHis tagged GFP:NFL head domain fusion protein. Unbound material was recovered and subjected to shotgun mass spectrometry in order to identify the hydrogel-bound proteins listed in Figure 1.

**Figure 1.**
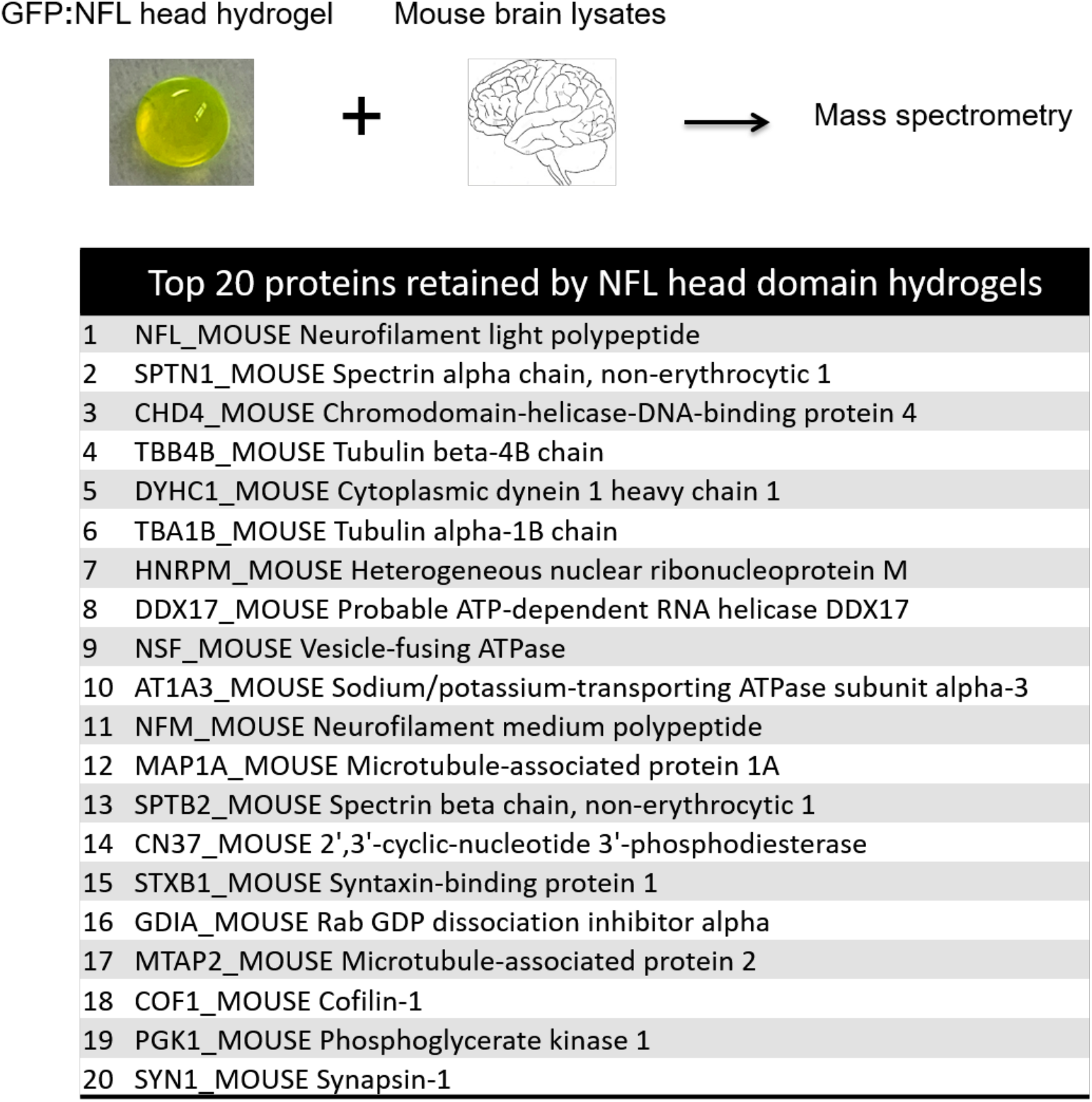
Binding of mouse brain proteins to hydrogel droplets formed from the NFL head domain. Hydrogel droplets were formed from a fusion protein linking 6xHis tagged GFP to an N-terminal fragment specifying the first 92 residues of the neurofilament light (NFL) protein (Experimental Procedures). Droplets 5 mm in size were formed on parafilm, incubated with a soluble lysate prepared from mouse brain tissue, washed and eluted by melting gel droplets in gelation buffer supplemented with 6 M guanidine-HCl. The melted GFP:NFL head domain protein was removed by Ni-affinity beads, allowing bound proteins to then be identified by shotgun mass spectrometry. The intact, endogenous NFL protein yielded a higher number of spectral counts than any other mouse brain protein bound to GFP:NFL hydrogel droplets. All peptides derived from the first 92 residues of the NFL protein were disregarded from the mass spectrometry data. Many other hydrogel-bound proteins are known from previous studies to interact with neuorfilaments.

### Evidence of specificity in intermediate filament head domain self-interaction

As deduced by quantitation of spectral counts, the top protein retrieved from mouse brain extracts by hydrogel droplets composed of NFL head domain polymers was the NFL protein itself. The majority of peptides identified in gel-bound samples corresponded to regions of NFL specifying its coiled-coil and tail domains. Since the hydrogel itself was prepared from a protein specifying the NFL head domain only (residues 2-92), it can be concluded that the NFL protein prominently bound by the hydrogel droplets was derived from the mouse brain extract.

Other than capturing the mouse brain NFL protein itself, NFL head domain hydrogel droplets selectively retained a number of axonal proteins including the α and β chains of non-erythrocyte spectrin, the α and β subunits of tubulin, three microtubule associated proteins designated MAP1A, MAP1B and MAP2, the heavy chain of cytoplasmic dynein, and the large plectin protein known to bridge association of assembled neurofilaments to microtubules (Figure 1). All nine of these proteins were already known to associate with axonal neurofilaments (Bocquet et al., 2009; Frappier et al., 1991; Ma et al., 2000; Miyata et al., 1986; Svitkina et al., 1996; Wagner et al., 2004; Yuan et al., 2017). Knowing that the medium and heavy neurofilament isoforms co-assemble with NFL (Yuan et al., 2017), we were not surprised to see NFM as the 11th protein on the list of mouse brain proteins bound to hydrogels formed from NFL head domain polymers, and NFH as the 111th protein on the list. Less expected was identification of two RNA-binding proteins, heterogeneous nuclear RNA ribonucleoprotein M (hnRNPM) and an ATP-dependent RNA helicase designated DDX17, as being prominently represented on the list of proteins bound by NFL head domain hydrogels.

Our simplistic interpretation of these observations is that the NFL head domain present in hydrogel droplets is able to specifically bind the head domain of the native NFL protein present in the soluble brain tissue lysate. This binding would be expected to secure retention of the entire NFL polypeptide, thus accounting for the abundant occurrence of tryptic peptides from the coiled-coil and tail domains of the protein. We further offer that the heterotypic trapping of other proteins known to interact with neurofilaments reflects the fact that these proteins may remain associated with NFL endogenous to the brain lysate through our conditions of binding, washing, and elution.

The aforementioned interpretation was tested using recombinant NFL protein samples linked to GFP. One fusion protein linked GFP to intact NFL, a second linked GFP to an NFL variant deleted of its head domain, and a third linked GFP to a variant deleted of the NFL tail domain. Each of these fusion proteins was expressed, purified, and incubated with hydrogel droplets composed of NFL head domain polymers. As shown in Figures 2A and 2B, NFL hydrogel droplets bound the GFP fusion proteins linked to either the intact NFL protein or the variant missing the unstructured tail domain. By contrast, the GFP fusion protein missing the unstructured NFL head domain was not trapped by hydrogels composed of the NFL head domain. As an extension of these experiments, GFP was fused to the isolated head, coiled-coil rod, or tail domains of NFL. Among the latter fusion proteins, only the GFP variant linked to the NFL head domain was retained by mCherry:NFL head domain hydrogel droplets (Figure 2B).

**Figure 2.**
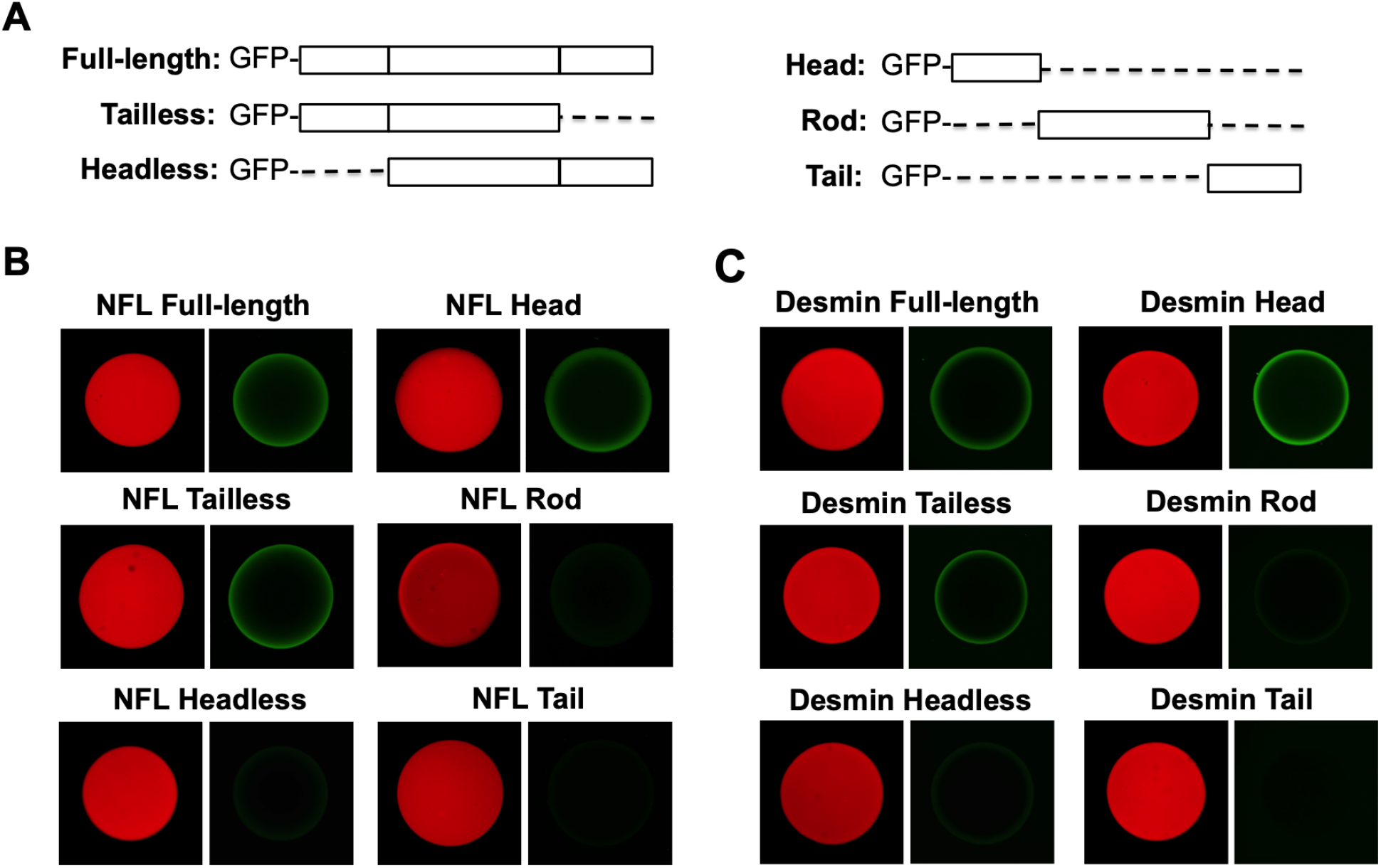
Hydrogel droplets formed from the NFL and desmin head domains bind cognate GFP-tagged proteins only if they retain the cognate head domain. Hydrogel droplets formed from fusion proteins linking mCherry to the head domains of NFL **(B)** or desmin **(C)** were exposed to GFP-tagged test proteins linked to the full-length versions of NFL or desmin, as well as versions missing the low complexity tail or head domain. GFP fusion proteins linked to the head domain alone, coiled-coil rod domain alone, or tail domain alone of either NFL or desmin were also tested for binding to hydrogel droplets composed of mCherry fused to the cognate head domain.

Similar experiments were conducted to study self-interaction of the head domain of desmin intermediate filaments. Desmin represents a canonical IF protein that has been extensively studied in the context of skeletal and cardiac muscle tissue (Paulin and Li, 2004). As shown in Figure 2C, hydrogel droplets composed of a fusion protein linking mCherry to the head domain of desmin were observed to bind a test protein that linked the intact desmin protein to GFP. A desmin variant deleted of its head domain failed to bind the hydrogel droplets, yet removal of the disordered tail domain of desmin did not impede gel binding. When fused in isolation to GFP, only the head domain of desmin allowed for GFP binding to mCherry:desmin head domain hydrogel droplets.

### ss-NMR spectra of NFL and desmin head domain polymers give evidence of structural order and β-strand secondary structure

Incubation of the isolated NFL and desmin head domains at a high protein concentration has been shown to yield amyloid-like polymers of uniform morphology (Lin et al., 2016). Unlike pathogenic, prion-like amyloids, polymers formed from the NFL or desmin head domains are labile to disassembly. Structural studies of these polymers were initiated by labeling the NFL and desmin head domains uniformly with ^13^C, allowing polymerization, and examining the samples by solid state nuclear magnetic resonance (ss-NMR) spectroscopy.

Figure 3 shows 2D ^13^C-^13^C spectra of NFL head domain polymers (Fig. 3A, left panel, blue contours) and desmin head domain polymers (Fig. 3B, left panel, blue contours), acquired at 13 °C as described in Experimental Procedures (see also Table S1). Crosspeak signals in these spectra are not sufficiently sharp to allow signals from individual residues to be resolved and assigned, but certain crosspeaks can be assigned to amino acid types, including Thr, Ser, Val, Pro, Ala, and Ile. With the exception of proline, the average ^13^C chemical shifts for these residues are consistent with a predominance of β-strand secondary structure in both NFL and desmin head domain polymers. This interpretation rests upon the observations that ^13^CO and ^13^C_α_ chemical shifts are smaller than random-coil values, and that ^13^C_β_ chemical shifts are larger than random-coil values (Table 1). The overall similarity of 2D ^13^C-^13^C ss-NMR spectra of the NFL and desmin head domain polymers reflects the similarity of their amino acid compositions (Supplemental Data, Figure S1).

**Figure 3.**
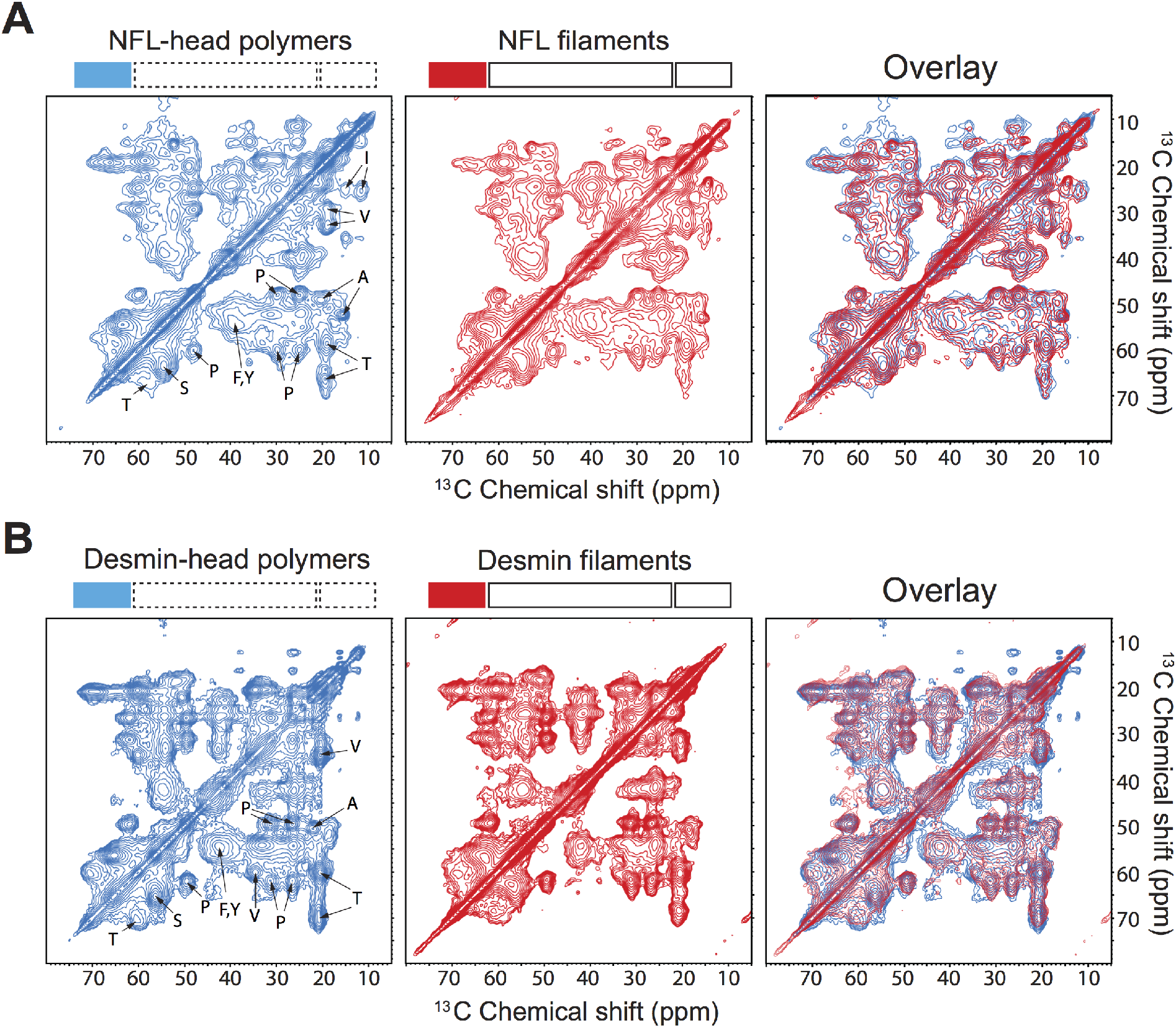
Two dimensional ^13^C-^13^C spectra of NFL and desmin head domain-only polymers as compared with segmentally-labeled NFL and desmin intermediate filaments. **Panel (A)** shows a 2D ^13^C-^13^C spectrum of ^13^C/^15^N uniformly-labeled NFL head domain-only polymers (left, blue) adjacent to a spectrum of the ^13^C/^15^N segmentally-labeled NFL head domain within assembled intermediate filaments (middle, red). Overlay of the two spectra reveal extensive overlap (right). Recording temperatures were 13 °C for head domain-only polymers and −23 °C for intermediate filaments. Plot counter increments are 1.4 for all spectra. **Panel (B)** shows a 2D ^13^C-^13^C spectra of ^13^C/^15^N uniformly-labeled desmin head domain-only polymers (left, blue) adjacent to a spectrum of the ^13^C/^15^N segmentally-labeled desmin head domain within assembled intermediate filaments (middle, red). Overlay of the two spectra reveals extensive overlap (right). Recording temperatures were 13 °C for the head domain-only polymers and − 26 °C for intermediate filaments. Plot counter increments are 1.4 for all spectra.

**Table 1:** Observed chemical shifts of several amino acids in the head domain polymers.

**Desmin Head Polymers**
| | C=O | Cα | Cβ |
| --- | --- | --- | --- |
| Pro | * | 62.1 (-1.2) | 31.7 (-0.4) |
| Val | 174.3 (-2.0) | 60.3 (-1.9) | 34.9 (+2.0) |
| Thr | 173.0 (-1.7) | 60.6 (-1.2) | 70.7 (+0.9) |
| Ser | 173.1 (-1.5) | 56.7 (-1.6) | 65.7 (+1.9) |
| Ala | 175.0 (-2.8) | 50.7 (-1.8) | 22.3 (+3.2) |
| Phe, Tyr | 173.9 (-2.0)<sup>a</sup> | 55.3 (-2.6)<sup>a</sup> | 41.8 (+3.0)<sup>a</sup> |

**NFL Head Polymers**
| | C=O | Cα | Cβ |
| --- | --- | --- | --- |
| Pro | * | 62.1 (-1.2) | 31.8 (-0.3) |
| Val | * | 60.6 (-1.6) | 35.0 (+2.1) |
| Thr | 172.4 (-2.3) | 60.3 (-1.5) | 70.0(+0.2) |
| Ser | 172.6 (-2.0) | 56.6 (-1.7) | 65.6 (+1.8) |
| Ala Peak1 | 174.5 (-3.3) | 51.2 (-1.3) | 22.2 (+3.1) |
| Peak2 | 178.6 (+0.8) | 54.4 (+1.9) | 17.5 (-1.6) |
| Phe, Tyr | 173.0 (-2.9)<sup>a</sup> | 56.5 (-1.4)<sup>a</sup> | 41.0 (+2.2)<sup>a</sup> |

The unit is point per million (ppm). The numbers in parentheses are differences from the chemical shifts for each amino acid in a random coil structure (Wishart et al., 1995).
\*peak not resolved. <sup>a</sup>The differences are calculated from the random coil chemical shifts of Tyr.

The observation that crosspeak signals from individual residues cannot be resolved in the 2D ss-NMR spectra of head domain polymers suggests that the level of structural order within these polymers is lower than in cross-β polymers formed by the low-complexity domains of FUS and hnRNPA2 (Murray et al., 2017; Murray et al., 2018) or in certain pathogenic amyloid fibrils (Colvin et al., 2016; Tuttle et al., 2016; Van Melckebeke et al., 2010), where better-resolved crosspeaks were observed. The precise level of order in NFL and desmin head domain polymers cannot be determined. However, it is notable that ss-NMR spectra of well-structured peptides and proteins in frozen solutions typically show ^13^C linewidths in the 1-3 ppm range (Hu et al., 2010; Jeon et al., 2019), attributable to small-amplitude fluctuations around an average structure that are not averaged out by thermal motions. Similar structural variations among protein molecules within the head domain polymers, corresponding to a root-mean-squared deviation of the atomic coordinates on the order of 2-4 Å, may account for the observed level of spectral resolution.

Most amyloid-like, cross-β protein assemblies contain in-register parallel β-sheets (Colvin et al., 2016; Murray et al., 2017; Murray et al., 2018; Tuttle et al., 2016). Exceptions to this rule include β-solenoidal fibrils (Van Melckebeke et al., 2010), worm-like metastable “protofibrils” (Qiang et al., 2012), and fibrils formed by certain short peptides (Petkova et al., 2004). To test for in-register parallel β-sheets in NFL and desmin head domain polymers, we prepared samples with ^13^C labels only at backbone carbonyl sites of Val, Leu, or Phe residues and used the PITHIRDS-CT ss-NMR technique to quantitatively measure ^13^C-^13^C dipole-dipole couplings (which are inversely proportional to the cube of internuclear distances). The PITHIRDS-CT data (Figure S2) indicate ^13^C-^13^C distances greater than 6 Å, inconsistent with in-register parallel β-sheets, for which ^13^C-^13^C distances of 4.8 Å are expected.

### NFL and desmin head domains display evidence of molecular structure in their respective, assembled intermediate filaments

Having observed that the isolated NFL and desmin head domains adopt a preferred molecular structure upon polymerization, the question arose as to whether this structure might be of biological relevance. If so, one would anticipate formation of much the same structure in fully mature NFL intermediate filaments. Recall that the coiled-coil domains of intermediate filaments become paired at an early step in filament assembly. This coiled-coil pairing positions two head domains in close proximity. Subsequent assembly of tetramers, elongated proto-filaments, and fully mature intermediate filaments positions 32 head domains in immediate, cylindrical proximity. These head domains ring the circumference of mature filaments and are disposed at roughly 45 nm intervals along the axial filament length (Alberts, 2002). The local concentration of NFL and desmin head domains must be high in these circumferential rings, perhaps sufficiently high to facilitate the weak, cross-β interactions formed when isolated head domains are incubated at a sufficiently high concentration to affect polymerization.

To test whether head domains adopt structures or structural distributions within assembled IFs that are similar to structures in head domain-only polymers, we acquired 2D ^13^C-^13^C spectra of segmentally labeled IFs (formed by full-length desmin or NFL, but with ^13^C-labeling localized only within the respective head domains via intein chemistry) (Figure S3). These 2D spectra are shown with red contours in the middle panels of Figure 3, and are superimposed upon the spectra of head domain-only polymers in the right panels. Crosspeak signal patterns of head domain polymers and segmentally labeled IFs are strikingly similar, providing strong evidence for structural similarity.

Although crosspeaks of individual residues are not resolved, the 2D ss-NMR spectra of head domain polymers and assembled IFs are strongly indicative of the presence of structural order. To probe this interpretation more rigorously, we prepared head domain-only polymers with ^13^C-^15^N labeling restricted to either Val and Ile residues (NFL) or Val and Thr (desmin). The spectra of these polymers was compared with the same protein samples dissolved in trifluoroacetic acid precipitated by cold ether. The latter treatment yielded amorphous aggregates not expected to exist in a structurally ordered state. 2D ^13^C-^13^C spectra of the Val/Ile (NFL) or Val/Thr (desmin) labeled head domain polymers and ether precipitates are shown in Figure 4. In these spectra, crosspeak signals of head domain polymers are clearly shifted away from the corresponding signals of amorphous head domain precipitates. The directions of shift correspond to expectations of a transition from a predominantly random coil structure in the amorphous material (red vertical lines in Figure 4) to a β-strand-enriched structure in the head domain polymers (blue vertical lines).

**Figure 4.**
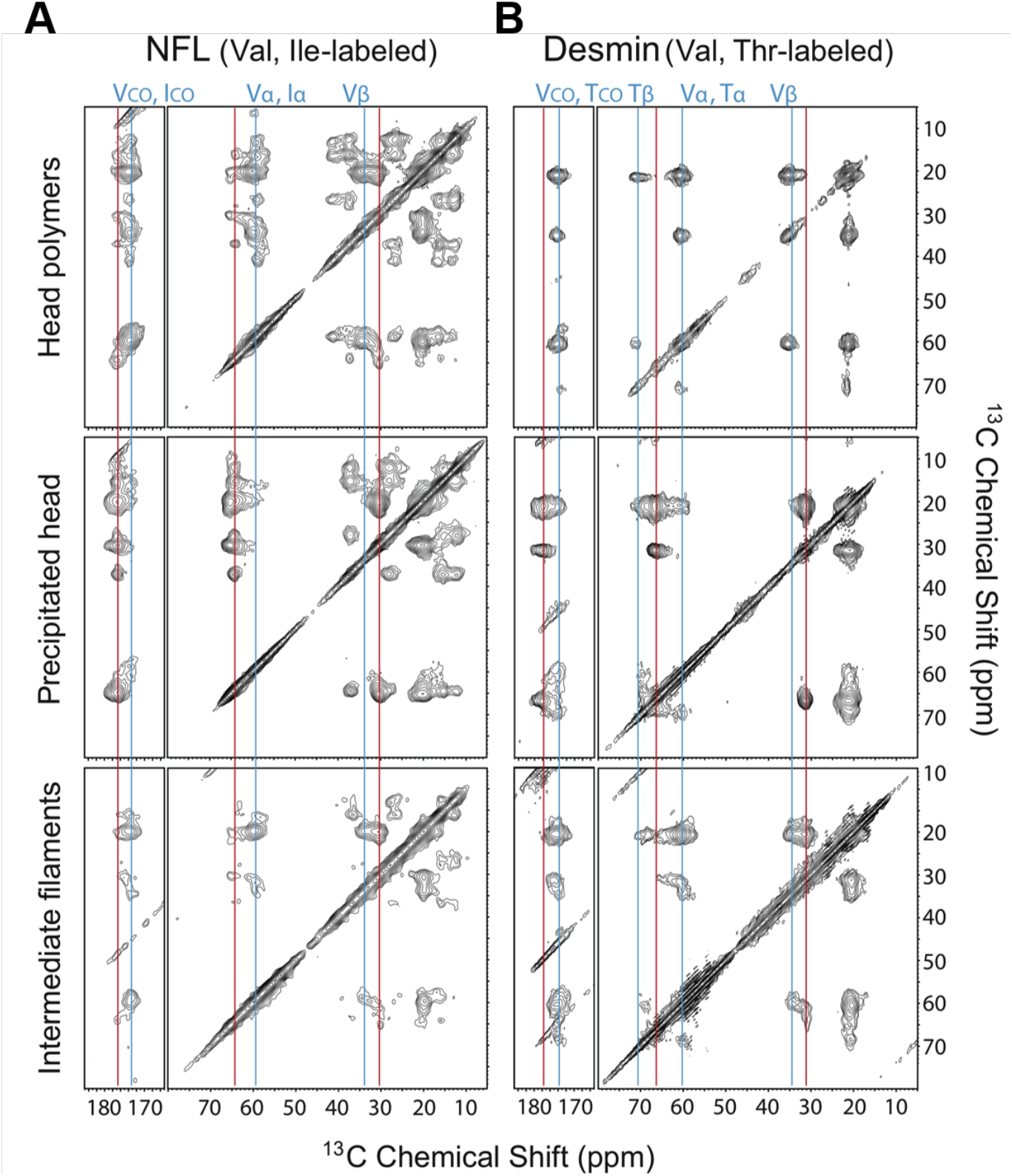
Two dimensional ^13^C-^13^C spectra of head domain-only polymers compared with assembled intermediate filaments segmentally labeled with specific amino acids. **Panel (A)** shows 2D ^13^C-^13^C spectra of ^13^C/^15^N Val and Ile-labeled NFL head domain-only polymers (top), an ether-denatured sample of the ^13^C/^15^N Val and Ile-labeled NFL head domain alone protein (middle), and ^13^C/^15^N Val and Ile segmentally-labeled NFL head domain within assembled intermediate filaments (bottom). Recording temperatures were 13 °C for the head domain-only samples and −23 °C for assembled intermediate filaments. Plot counter increments are 1.4 for all spectra. **Panel (B)** shows 2D ^13^C-^13^C spectra of ^13^C/^15^N Val and Thr-labeled desmin head domain-only polymers (top), an ether-denatured sample of the ^13^C/^15^N Val and Thr-labeled desmin head domain alone protein (middle), and ^13^C/^15^N Val and Thr segmentally-labeled desmin head domain within assembled intermediate filaments (bottom). Recording temperatures were 13 °C for the head domain-only samples and −26 °C for assembled intermediate filaments. Plot counter increments are 1.4 for all spectra. Vertical blue lines indicate peak positions for β-strand-like crosspeak signals in spectra of head domain-only polymers. Vertical red lines indicate non-β-strand-like signals in spectra of ether-precipitated samples.

Intein-mediated chemistry was used to introduce the same Val/Ile isotopes into the head domain of the full-length NFL, and the same Val/Thr isotopes into the head domain of the full-length desmin. Following purification, the segmentally labeled samples were assembled into mature intermediate filaments and analyzed by ss-NMR. The same patterns of crosspeak signal shift were observed in the assembled NFL and desmin intermediate filaments as had been seen in the head domain-only polymers. Although not all Val/Ile (NFL) or Val/Thr (desmin) crosspeak signal intensity in 2D spectra of the assembled IFs aligned exactly with signal intensity of 2D spectra of the head domain-alone polymers, spectra of the Val/Ile- and Val/Thr-labeled IFs clearly agree more closely with spectra of head domain polymers than with spectra of amorphous, ether-precipitated material.

### NFL and desmin head domains are partially dynamic in intermediate filaments at biologically relevant temperatures

Two dimensional ss-NMR spectra of IFs in Figures 3 and 4 were recorded at low temperatures (−26 °C for desmin, −23 °C for NFL) because spectra recorded above 0 °C showed significantly weaker crosspeaks (Figure S4). The reductions in crosspeak intensities at higher temperatures are attributable to molecular motions that partially average out nuclear magnetic dipole-dipole couplings, which drive the nuclear spin polarization transfers that produce crosspeaks. Apparently, the molecular structures and degree of conformational disorder of head domains in head domain-only polymers and IFs are similar, but the head domains are more highly dynamic (*i.e*., “mushy”) in IFs above 0 °C. Conformational variations in IFs are not necessarily larger in amplitude at higher temperatures, but occur on shorter time scales. Once the motional time scales exceed roughly 1 ms, motional effects on the 2D ^13^C-^13^C ss-NMR spectra become minimal.

To probe molecular motions of head domains in the IFs in more detail, we recorded 1D ^13^C NMR spectra of the segmentally ^13^C-labeled IFs with two different methods. To observe signals from relatively rigid protein segments, we used ^1^H-^13^C cross-polarization (CP) (Pines et al., 1973), driven by nuclear magnetic dipole-dipole couplings, to prepare transverse ^13^C polarization and high-power ^1^H decoupling with two-pulse phase modulation (TPPM) (Bennett et al., 1995) during signal acquisition. To observe signals from highly dynamic segments with nearly isotropic motions, we used the “insensitive nuclei enhanced by polarization transfer” (INEPT) (Morris and Freeman, 1979) technique, driven by scalar couplings, to prepare transverse ^13^C polarization and low-power composite pulse decoupling (Levitt and Freeman, 1981; Tycko et al., 1985). As shown in Figure 5 for both NFL IFs and desmin IFs, CP-based signals increase in intensity with decreasing temperature, while INEPT-based signals decrease in intensity.

**Figure 5.**
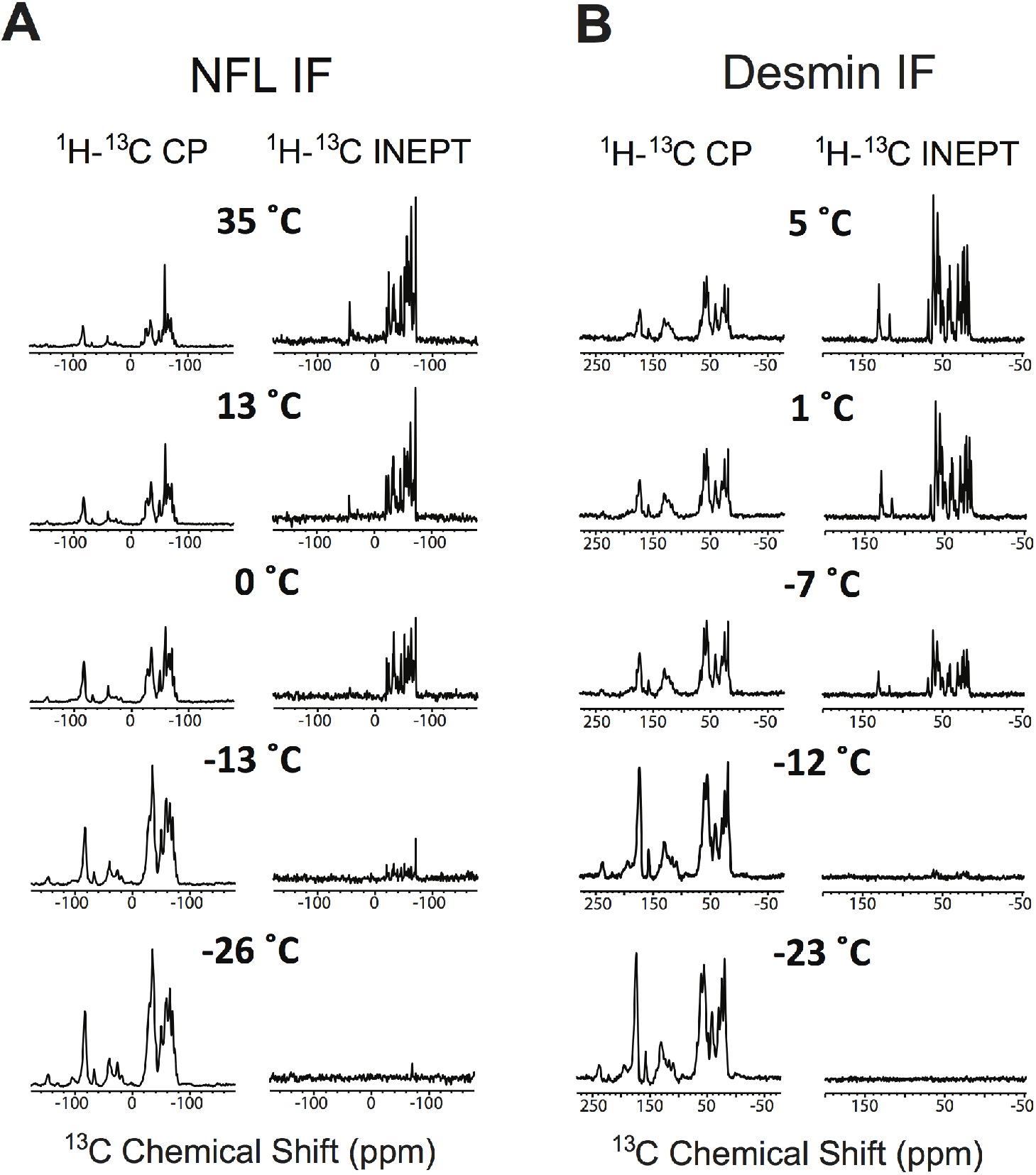
1D cross polarization and INEPT spectra of segmentally-labeled intermediate filaments. **Panel (A)** shows one dimensional ^1^H-^13^C cross polarization and INEPT spectra of C/ N segmentally-labeled NFL head domain as situated *in situ* in the context of assembled intermediate filaments. Spectra were recorded at temperatures ranging from 35 °C to −26 °C as indicated. **Panel (B)** shows one dimensional ^1^H-^13^C cross polarization and INEPT spectra of ^13^C/^15^N segmentally-labeled desmin head domain as situated *in situ* in the context of assembled intermediate filaments. Spectra were recorded at temperatures ranging from 5 °C to −23 °C as indicated. Spectra within each column are plotted on the same vertical scale.

We interpret the data in Figures 5 and S4 as evidence that both desmin and NFL head domains contain structured segments, with moderate-amplitude disorder as discussed above, and unstructured segments. The structured segments have limited internal motion, occurring on sub-millisecond time scales at the higher temperatures in these experiments and >1 ms time scales at the lowest temperatures. The unstructured segments have large-amplitude motions (*i.e*., are dynamically disordered) at the higher temperatures, on sub-microsecond time scales. Motions of the unstructured segments are reduced at lower temperatures, leading to inefficient INEPT polarization transfers and weak signals in INEPT-based spectra. Importantly, it is primarily the overall signal intensities, rather than the peak positions and line shapes, that change with temperature in both the CP-based and the INEPT-based spectra in Figure 5. Therefore, these spectra should not be interpreted to be giving evidence of a conversion of disordered segments to ordered segments with decreasing temperature.

### Phosphorylation of the NFL and desmin head domains disassembles IFs and inhibits head domain self-association

Intermediate filaments have long been known to disassemble during mitosis (Chou et al., 1990; Rosevear et al., 1990). In numerous instances disassembly has been attributed to phosphorylation of IF head domains (Cleverley et al., 1998; Geisler and Weber, 1988; Hisanaga et al., 1990; Hisanaga et al., 1994; Perrot et al., 2008; Winter et al., 2014; Yuan et al., 2017). Upon incubation of either NFL or desmin IFs with a combination of ATP and protein kinase A (PKA), both filaments were observed to disassemble (Figure 6). Exposure of either IF to either ATP or PKA alone failed to affect filament disassembly. The combination of ATP and PKA also effected release of the GFP:NFL head domain fusion protein from mCherry:NFL head domain hydrogel droplets. Likewise, the GFP:desmin head domain fusion protein bound to mCherry:desmin head domain hydrogel droplets was released upon exposure to both ATP and PKA (Figure 6). Reactions missing ATP or PKA left the respective GFP fusion proteins hydrogel-bound.

**Figure 6.**
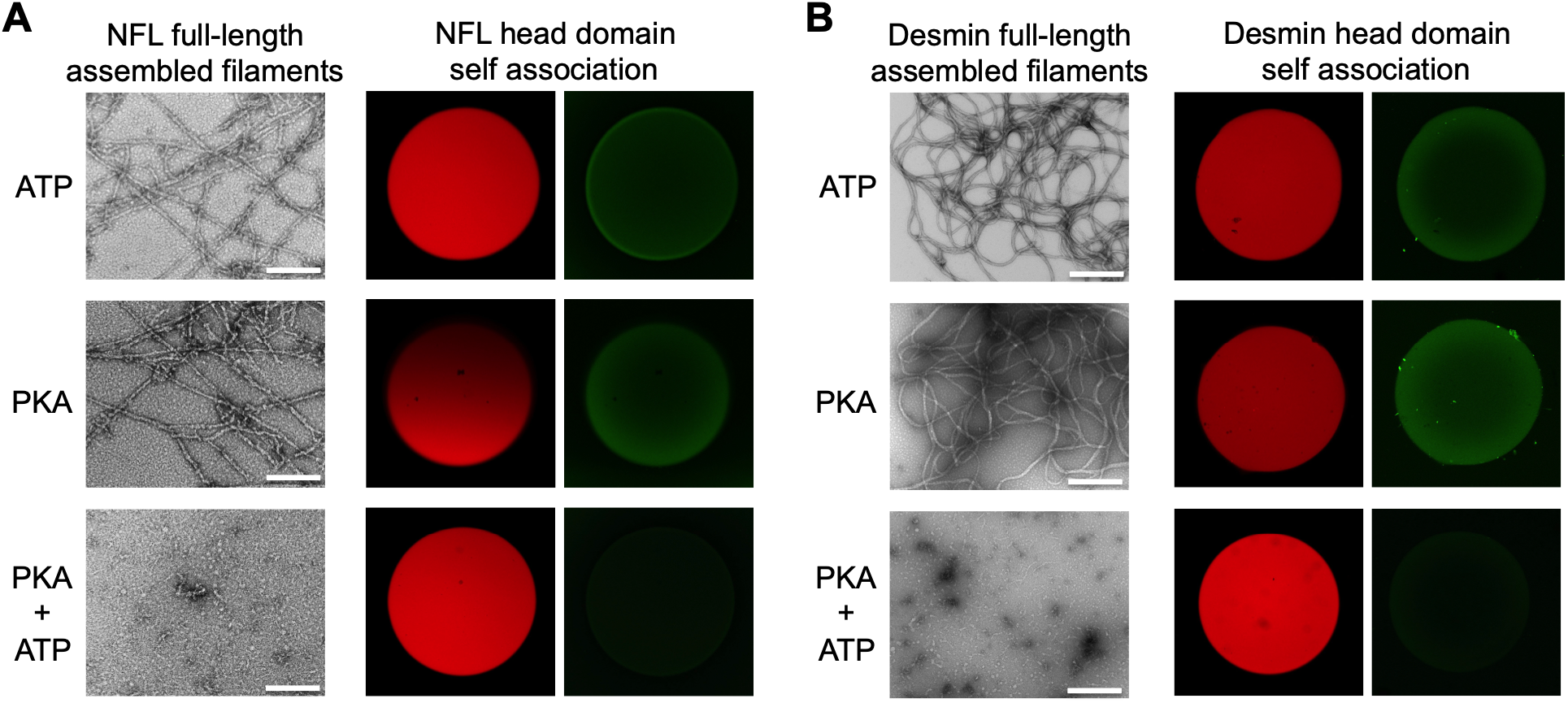
Protein kinase A-mediated disassembly of NFL and desmin intermediate filaments and release of GFP:head domain test proteins from cognate hydrogel binding. Assembled intermediate filaments prepared from full-length NFL and desmin were incubated with ATP alone, protein kinase A (PKA) alone, or both ATP and PKA. PKA-mediated phosphorylation led to disassembly of both types of intermediate filaments as deduced by transmission electron microscopy. Hydrogel droplets formed from mCherry fused to the head domain of either NFL **(A)** or desmin **(B)** were pre-bound with GFP fusion proteins linked to the head domain of either NFL or desmin. Samples were incubated with ATP alone (top images), PKA alone (middle images) or both agents (bottom images). Release of hydrogel bound GFP-tagged protein was only observed upon co-incubation with both ATP and PKA. All scale bars =200 nm.

### Recurrent mutations in NFL and desmin head domains accelerate β-strand-enriched self-association

Human genetic studies have identified perplexingly recurrent, autosomal dominant mutations localized to the NFL head domain. Studies of pedigrees tracing the genetic cause of neurological deficits, commonly described in the context of Charcot-Marie-Tooth (CMT) disease, have identified mutations in one of two proline residues located within the NFL head domain (Jordanova et al., 2003; Shin et al., 2008). Independent familial mutations changing proline residue 8 to leucine, arginine, or glutamine are commonly understood to be causative of CMT disease. Surprisingly, independent familial mutations causative of CMT disease have also been found to change proline 22 within the NFL head domain to serine, threonine, or arginine. The reason why CMT disease-causing mutations are recurrently found in these two proline residues has been obscured by the assumption that the head domains of all intermediate filament proteins function in the absence of molecular order. If a protein segment is intrinsically disordered, why would its biological function care about the subtle change of but a single amino acid?

NFL variants carrying a P-to-Q, P-to-L, or P-to-R mutation at residue 8, or a P-to-S, P-to-T, or P-to-R mutation at residue 22, were expressed as recombinant protein, purified, and tested for the formation of intermediate filaments. All six mutations significantly impeded IF assembly, yielding much shorter filaments relative to the native NFL protein (Figure 7A and S5A) (Sasaki et al., 2006). In order to study these NFL variants more carefully, head domains bearing three different P8 mutations, and three different P22 mutations, were expressed in bacterial cells, purified, and assayed for their capacity to form labile, cross-β polymers. Our standard methods involve purification of the NFL head domain in 8M urea followed by sequential dialysis into gelation buffer supplemented by 4 M, 2 M, 1 M, 0.5 M and no urea (Experimental Procedures). Upon initiating studies of the six different CMT-causing mutations, it was noticed that the mutated variants became cloudy upon dialysis of the protein samples from 8 M to 4 M urea (Figure 7C). Analysis of the cloudy samples by transmission electron microscopy revealed uniform, amyloid like polymers indistinguishable from the labile, cross-β polymers formed by the native NFL head domain (Figure 7C and S5B) which do not form without complete removal of the urea denaturant.

**Figure 7.**
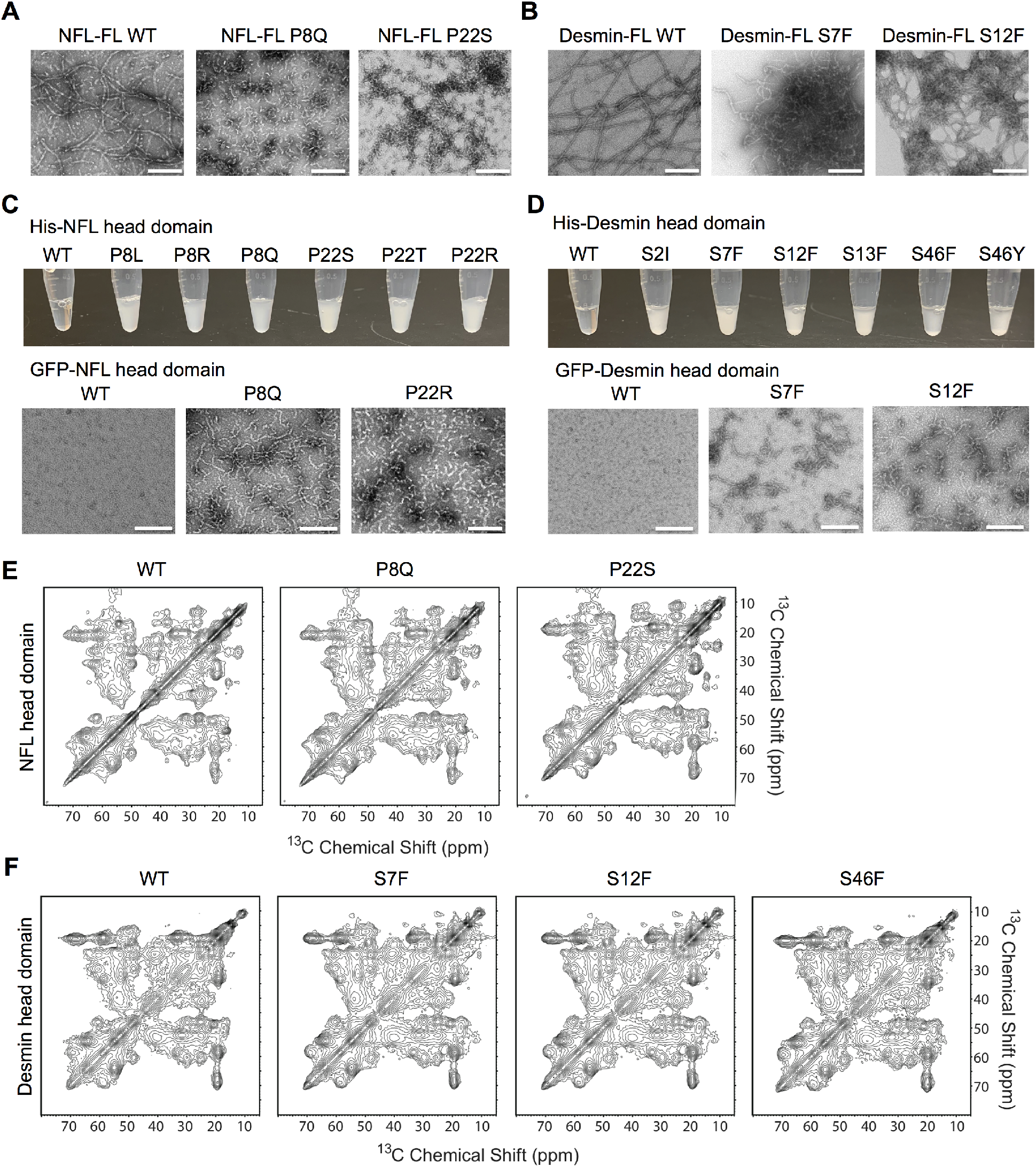
Effects of disease-causing mutations within NFL and desmin head domains on assembly of intermediate filaments (A and B), cross-β polymerization of isolated head domains (C and D), and ss-NRM spectra of isotopically labeled head domains (E and F). Full-length (FL) NFL and desmin proteins bearing indicated head domain mutations were incubated under conditions suitable for assembly of intermediate filaments. Compared with native (WT) proteins, disease-causing mutations in the NFL and desmin head domains led to the formation of either short (NFL) or tangled (desmin) intermediate filaments (**panels A and B**, and Figure S5A). Isolated head domain samples of native (WT) or six different NFL head domain mutants were tested for formation of cross-β polymers **(C)**. Relative to the native NFL head domain, all six head domain mutants formed polymers prematurely in the presence of 4 M urea as revealed by opalescence. Transmission electron microscopy showed no polymers formed from the native NFL head domain in urea (experimental procedures), but clear evidence of polymers formed from the P8Q and P22R head domain mutants. Isolated head domain samples of native (WT) and six desmin head domain mutants were also tested for formation of cross-β polymers in the presence of 3 M urea **(D)**. Relative to the native desmin head domain, all six head domain mutants formed polymers prematurely in the presence of 3 M urea. Transmission electron microscopy revealed no polymers formed from the native desmin head domain in urea (experimental procedures), but clear evidence of polymers formed from the S7F, S12F and S46F head domain mutants. Two-dimensional cross polarization ss-NMR spectra of isotopically labeled head domains of NFL and desmin showed no differences between native (WT) proteins and indicated head domain mutants **(E and F)**. All scale bars =200 nm.

To compare native and mutant polymers in greater detail, we isotopically labeled the P8Q and P22S CMT variants of the NFL head domain with ^13^C and ^15^N, allowed for polymerization, and examined the material by ss-NMR. As shown in Figure 7E, 2D ^13^C-^13^C ss-NMR spectra of native and mutant NFL head domain polymers were indistinguishable. We conclude from these observations that mutation of either proline residue 8 or proline residue 22 to any of a number of different amino acids enhances the ability of the NFL head domain to self-associate in the form β-strand-enriched polymers without grossly changing protein structure.

Human genetic studies have also led to the discovery of autosomal dominant mutations localized to the head domain of desmin. Pedigree studies have associated desmin head domain mutations with cardiomyopathy (Azzimato et al., 2016; Clemen et al., 2013). Surprisingly, all six of these desmin head domain mutations change individual serine residues, including S2I, S7F, S12F, S13F, S46F and S46Y. The latter five mutations change a single serine residue to either phenylalanine or tyrosine. All of these disease-causing desmin variants were expressed as recombinant protein, purified and tested for the formation of intermediate filaments. All six mutants impeded IF assembly yielding tangled and aggregated filaments (Figure 7B and S5A) (Sharma et al., 2009).

When expressed in the context of the isolated desmin head domain and assayed for formation of β-strand-enriched polymers, all six variant proteins, including S2I, S7F, S12F, S13F, S46F and S46Y, aberrantly polymerized in the presence of 3 M urea, just as was observed for the six different CMT-causing P8 and P22 mutations within the NFL head domain (Figures 7D and S5B). The S7F, S12F and S46F variants of the desmin head domain were labeled with ^13^C and ^15^N, allowed to polymerize, and evaluated by ss-NMR. As shown in Figure 7F, all three mutants exhibited 2D ^13^C-^13^C ss-NMR spectra indistinguishable from polymers made from the native desmin head domain.

## Discussion

Our interest in the assembly of intermediate filaments came by accident in studies of the aliphatic alcohol, 1,6-hexanediol. This agent had long been known to compromise the permeability barrier of nucleopores and melt nuclear and cytoplasmic puncta not surrounded by investing membranes (Kroschwald et al., 2015; Patel et al., 2007; Updike et al., 2011). We and others had noticed that 1,6-hexanediol also melts phase separated hydrogels and liquid-like droplets formed in test tubes via self-association of low complexity domains (Lin et al., 2016; Molliex et al., 2015). As a control experiment, we asked whether 1,6-hexanediol might melt cellular structures in a specific or non-specific manner. It was observed that, upon administration to cultured mammalian cells, the agent did not disassemble either actin filaments or microtubules. Much to our surprise, however, 1,6-hexanediol caused dramatic disassembly of intermediate filaments specified by either vimentin or keratin (Lin et al., 2016). We quickly made note of extensive literature showing that intermediate filament assembly is reliant upon head domains universally specified by low complexity sequences, and proceeded to confirm that the head domains of six different intermediate filament proteins display the ability to become phase separated in a manner indistinguishable from prototypic low complexity domains associated with RNA-binding proteins.

Here we described experiments focused upon the amino-terminal head domains of the NFL and desmin intermediate filament proteins. Both head domain are of low sequence complexity and have long been understood to function in a state of intrinsic disorder (Herrmann and Aebi, 2016). Evidence is shown that the NFL and desmin head domains, when studied in isolation, can form labile β-strand-enriched polymers (Lin et al., 2016). Hydrogel preparations composed of NFL head domain polymers, upon incubation with total soluble lysate from mouse brain tissue, selectively bind the endogenous NFL protein (Figure 1). Capture of the intact NFL protein in this hydrogel binding reaction was shown to be dependent upon the presence of the NFL head domain (Figure 2). Similarly conceived experiments focused on the desmin head domain yielded similar observations. We conclude from these observations that the head domains of NFL and desmin are capable of some means of self-association that is of demonstrably significant specificity.

In addition to the selective capture of the mouse brain NFL protein, hydrogel droplets composed of the NFL head domain also captured a number of neuronal proteins known to associate with neurofilaments, including tubulin, spectrin, and microtubule associated proteins (MAPs) (Figure 1). Surprisingly, two RNA-binding proteins were also captured by NFL hydrogels, hnRNPM and DDX17. Histological studies of cultured nerve cells have revealed hnRNPM at synaptic terminals, giving evidence that this RNA-binding protein may participate in the control of localized, synaptic translation (Zhang et al., 2012). We offer that interaction between RNA granules, perhaps containing hnRNPM, and neurofilaments may assist in the transport of mRNAs to specialized regions of neurons for the purpose of localized translation.

Previously reported electron microscopic studies have revealed binding of a GFP protein linked to the low complexity domain of the fused in sarcoma (FUS) RNA-binding protein to vimentin intermediate filaments (Lin et al., 2016). The reported binding was observed at 45 nm intervals along the axial length of vimentin filaments, corresponding to the repetitive, circumferential organization of the vimentin head domain as it helps organize filament architecture. Knowing that RNA granules in the eggs of both flies and frogs co-localize with intermediate filaments (Gaspar et al., 2017; Pondel and King, 1988), the observations reported herein may be offering mechanistic insight relevant to how cytoplasmic RNA granules come to be localized in eukaryotic cells.

Solid state NMR methods were used to ask whether the NFL and desmin head domains might employ molecular structure to facilitate the specificity of self-association. Our studies give evidence of β-strand-enriched interactions that define the structural interface dictating specificity of NFL and desmin head domain self-association (Table 1, Figure 3 and 4). By use of intein chemistry facilitating segmental labeling, intact NFL and desmin polypeptides were produced bearing several different forms of isotope labeling restricted to the respective head domains of the two proteins. Subsequent to assembly into mature intermediate filaments, the isotopically-labeled head domains of NFL and desmin were evaluated by ss-NMR. As a function of sequential cooling, we observed the appearance of cross-polarization spectra highly similar to spectra found in polymeric samples formed from the NFL and desmin head domains alone (Figure 3 and 4). From these observations we offer three conclusions. First, we propose that the observed structural interactions are responsible for the specificity of NFL and desmin head domain self-interaction. Second, we propose that these interactions are responsible for the augmentative role of the respective head domains in assembly of mature NFL and desmin intermediate filaments. Third, we propose that the observed structural interactions are inherently dynamic – with their diagnostic NMR spectra becoming enhanced upon cooling (Figure 5 and S4).

Self-associative interactions specified by the NFL and desmin head domains appear, by evolutionary design, to be inherently weak and readily disrupted by phosphorylation (Figure 6). We proposed that these head domains, even in the context of assembled IFs, are continuously moving in and out of the structurally ordered state. If so, access to enzymes affecting post-translational modification might be facile. Protein kinase A (PKA) has been reported to phosphorylate the NFL head domain on six different serine residues, including S2, S12, S41, S49, S55 and S62 (Cleverley et al., 1998). The desmin head domain is reported to be phosphorylated by PKA on residues S45 and S60, protein kinase C on S13, S48 and S68, and cyclin-dependent kinase on S7, S32 and T76 (Winter et al., 2014). We as yet do not know the location of these sites with respect to the β-strand-enriched structural regions of either the NFL or desmin head domains. If proximal to transiently structured regions, phosphate groups might impart repulsive charge:charge interactions incompatible with the structured state as has been observed for DNA-dependent protein kinase-mediated phosphorylation and disruption of cross-β structural interactions formed by the fused in sarcoma (FUS) LC domain (Murray et al., 2017).

Quite by contrast to the hypothetically destabilizing influences of head domain phosphorylation, we conclude that disease causing mutations in the NFL and desmin head domains enhance self-association (Figure 7). All such mutations are autosomal dominant (Clemen et al., 2013; Jordanova et al., 2003; Shin et al., 2008), and are most easily understood to elicit gain-of-function phenotypic effects. CMT-causing mutations in the NFL gene recurrently localize to proline residues 8 or 22. All such mutations were observed to enhance head domain self-association (Figure 7 and S5). Paradoxically, enhancement of NFL head domain self-interaction was observed to be detrimental to proper assembly of neurofilaments. It was similarly observed that recurrent, disease-causing mutations in the desmin head domain also enhance head domain self-association and impede IF assembly (Figure 7 and S5). At present, we remain ignorant as to why enhanced self-association of NFL and desmin head domains interferes with IF assembly.

The studies reported herein combine human genetics, biochemistry and NMR spectroscopy to study the head domains of NFL and desmin. As such, this work is reminiscent of similarly interwoven studies on RNA-binding proteins. Idiosyncratically recurrent mutations of an analogous aspartic acid residue present in the low complexity domains of the hnRNPA1, hnRNPA2 and hnRNPDL proteins lead to various forms of neurologic disease (Kim et al., 2013; Vieira et al., 2014). The relevant aspartic acid residue of the three hnRNP proteins is localized within self-associating cross-β regions that allow for formation of both liquid-like droplets and hydrogel polymers. It has been interpreted that these aspartic acid residues impart repulsive, charge:charge interactions that are structure-destabilizing. Upon mutational change, often times to valine, the destabilizing interactions are removed, leading to enhanced self-association of the hnRNP LC domains and age-related neurodegenerative disease (Gui et al., 2019; Murray et al., 2018). We offer that human genetic studies are helping us to understand that evolution has established a proper balance of “designed instability” to self-associative, β-strand-enriched interactions that are of biologic utility to both RNA-binding proteins and intermediate filament proteins.

Five of the six disease-causing mutations in the desmin head domain change individual serine residues to either phenylalanine or tyrosine (S7F, S12F, S13F, S46F and S46Y). The importance of aromatic amino acids to LC domain self-association was proposed and functionally confirmed in the earliest studies of phase transition of both the FG repeats of nucleoporins (Frey and Gorlich, 2007) and the LC domain of the fused in sarcoma (FUS) RNA-binding protein (Han et al., 2012; Kato et al., 2012). We speculate that the substitution of hydrophilic serine residues by either phenylalanine or tyrosine somehow accelerates or enhances desmin head domain self-association. At the same time, we appreciate the possibility that S-to-F or S-to-Y mutations might eliminate sites of phosphorylation important to the regulation of desmin assembly in living cells.

We close by asking why proline residues 8 and 22 are so important for proper function of the NFL head domain? CMT-causing mutations in the NFL protein are restricted to P8 and P22, and disease can result from mutations that change either of these proline residues to any of a number of different amino acids. Proline is an unusual amino acid residue for two reasons. First, it is an imino acid extending a covalently closed ring of three CH2 groups extending off the peptide nitrogen and back onto the adjacent α-carbon to which the side chains of the other 19 amino acids are invariably attached. This chemical conformation eliminates the opportunity for the peptide backbone of proline residues to participate in the hydrogen bonding network essential for β-strand interactions (Pauling and Corey, 1951). Second, the peptide bond adjacent to proline residues can rotate between the *cis* and *trans* states. The peptide bond adjacent to all other amino acids specifying biological polypeptides is invariably restricted to the *trans* conformation. Mutation of proline to any of the other 19 amino acids would – at the same time - restore the opportunity for β-strand hydrogen bonding and confine the adjacent peptide bond to the *trans* state. Deeper experimental investigation into the structure and function of the NFL head domain may allow us to probe these perplexing observations.

## Acknowledgments

We thank Yonghao Yu for extensive help with mass spectrometry, Deepak Nijhawan for thoughtful advice on human genetic mutations in the NFL and desmin head domains, and the Electron Microscopy Core Facility of UT Southwestern Medical Center for technical support. This work was supported by grant 5R35GM13130358 from the NIGMS and unrestricted funding provided to SLM by an anonymous donor.

## Supplementary figures and table

**Figure S1.**
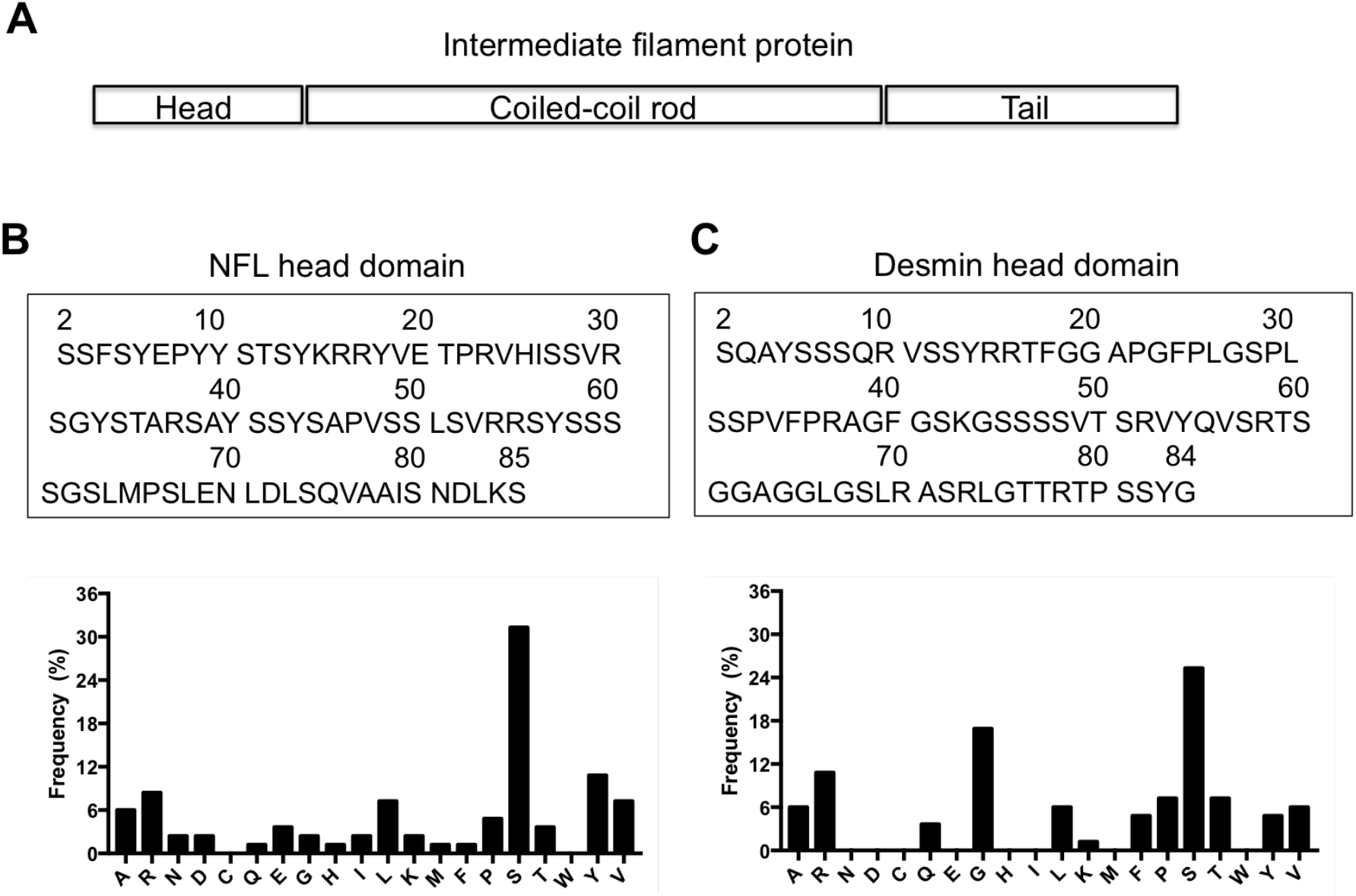
Amino acid sequences and composition of NFL and desmin head domains. (**A**) Schematic diagram of intermediate filament domain architecture. **(B)** amino acid sequence (above) and composition (below) of NFL head domain. **(C)** amino acid sequence (above) and composition (below) of desmin head domain.

**Figure S2.**
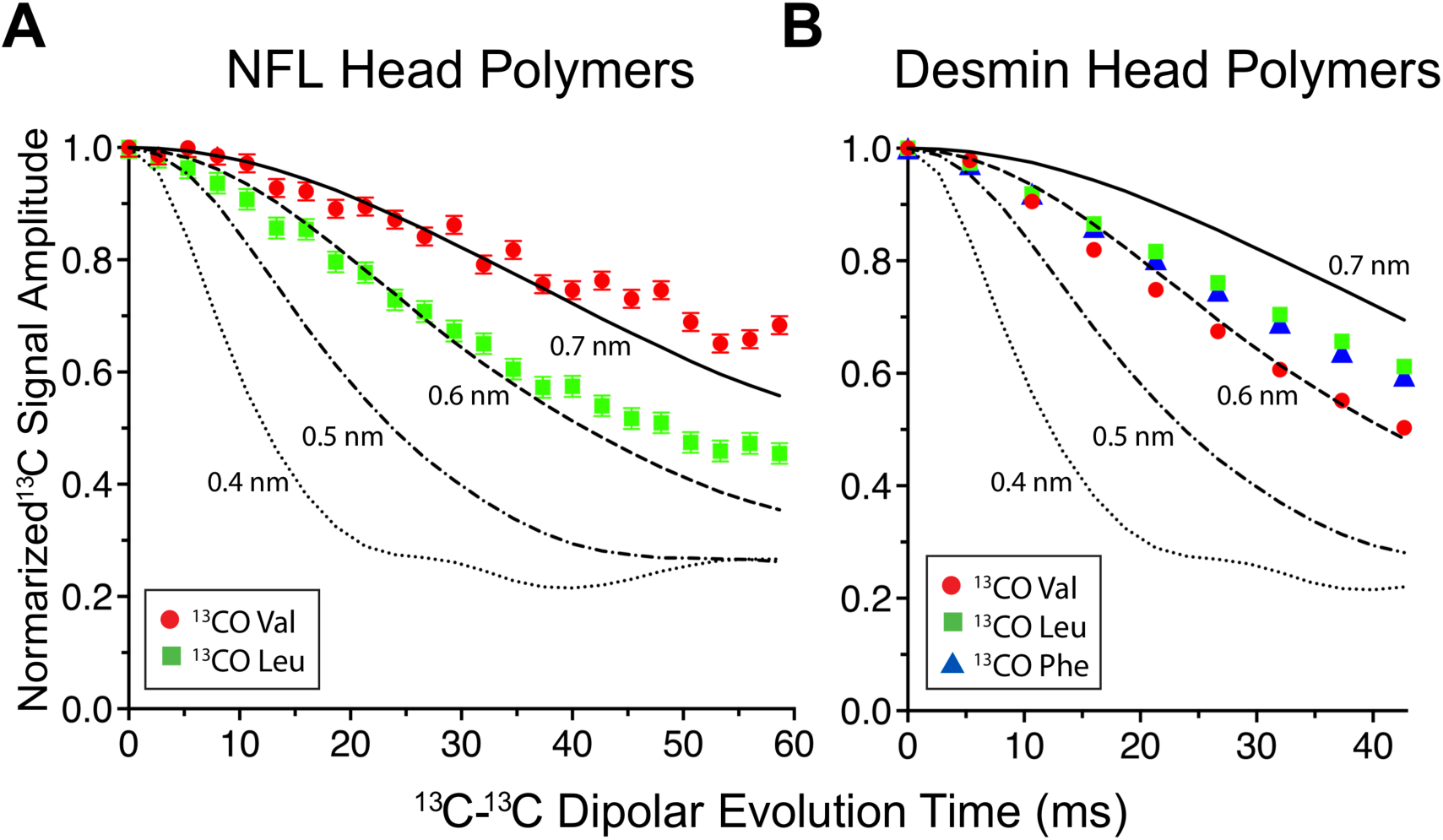
Quantitative intermolecular distance measurements by the ^13^C-^13^C dipole-dipole recoupling technique. **(A)** ^13^C-^13^C PITHIRDS-CT dipolar recoupling data for NFL head domain-only polymers that were ^13^C-labeled only at backbone carbonyl sites of all Val or Leu residues within the head domain. Normalized peak intensities at different dipolar evolution times are plotted. **(B)** ^13^C-^13^C PITHIRDS-CT dipolar recoupling data for desmin head domain-only polymers that were ^13^C-labeled only at backbone carbonyl sites of all Val, Leu or Phe residues within the head domain. Normalized peak intensities at different dipolar evolution time are plotted. Lines represent simulated data for linear chains of ^13^C label with the indicated interatomic spacings.

**Figure S3.**
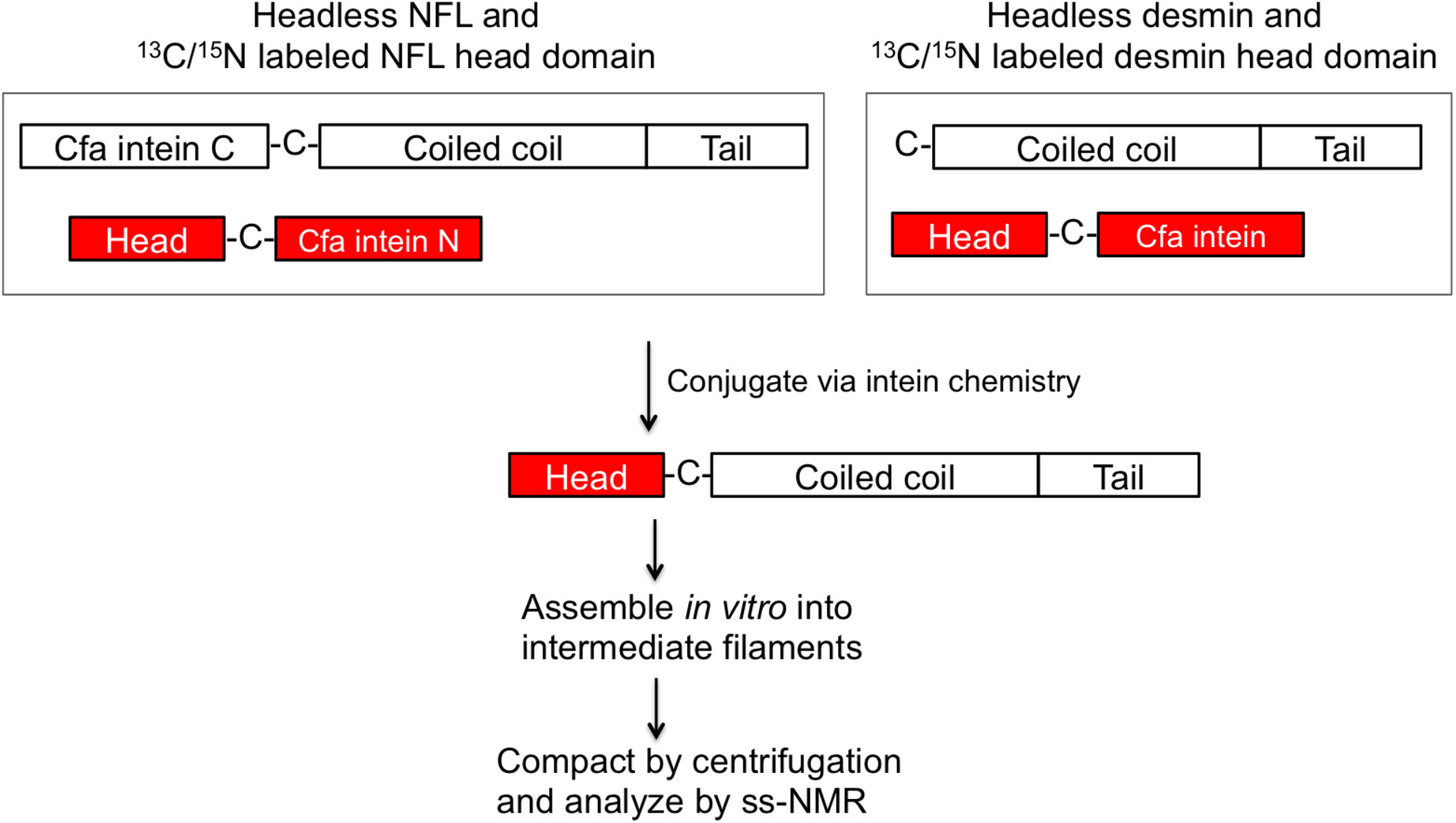
Scheme of intein chemistry used for segmentally labeling of NFL and desmin full-length proteins with isotopically labeled head domains. Split intein chemistry was applied for NFL head domain labeling (**top left panel**). Unlabeled headless NFL was N-terminally fused with C-terminal half of Cfa intein (Cfa intein C). Isotopically labeled NFL head domain was C-terminally fused with N-terminal half of Cfa intein (Cfa intein N). Unlabeled, headless NFL derivative, and isotope-labeled head domain derivative were expressed as N-terminal GFP tagged proteins and digested by caspase 3 at designed site to remove the GFP tag (Experimental Procedures) and conjugated via split intein reaction. An intact intein reaction was applied for segmental labeling of the desmin head domain (**top right panel**). Isotopically-labeled desmin head domain was C-terminally fused to an intact Cfa intein. Both isotopically-labeled desmin head domain and unlabeled headless desmin were expressed as N-terminal His tagged proteins and digested with caspase 3 to remove the His tag (Experimental Procedures). Isotopically-labeled desmin head domain and unlabeled headless desmin were conjugated via native chemical ligation reaction. Full-length products of NFL and desmin were assembled into filaments and compacted by ultracentrifugation for analysis by solid state NMR spectroscopy.

**Figure S4.**
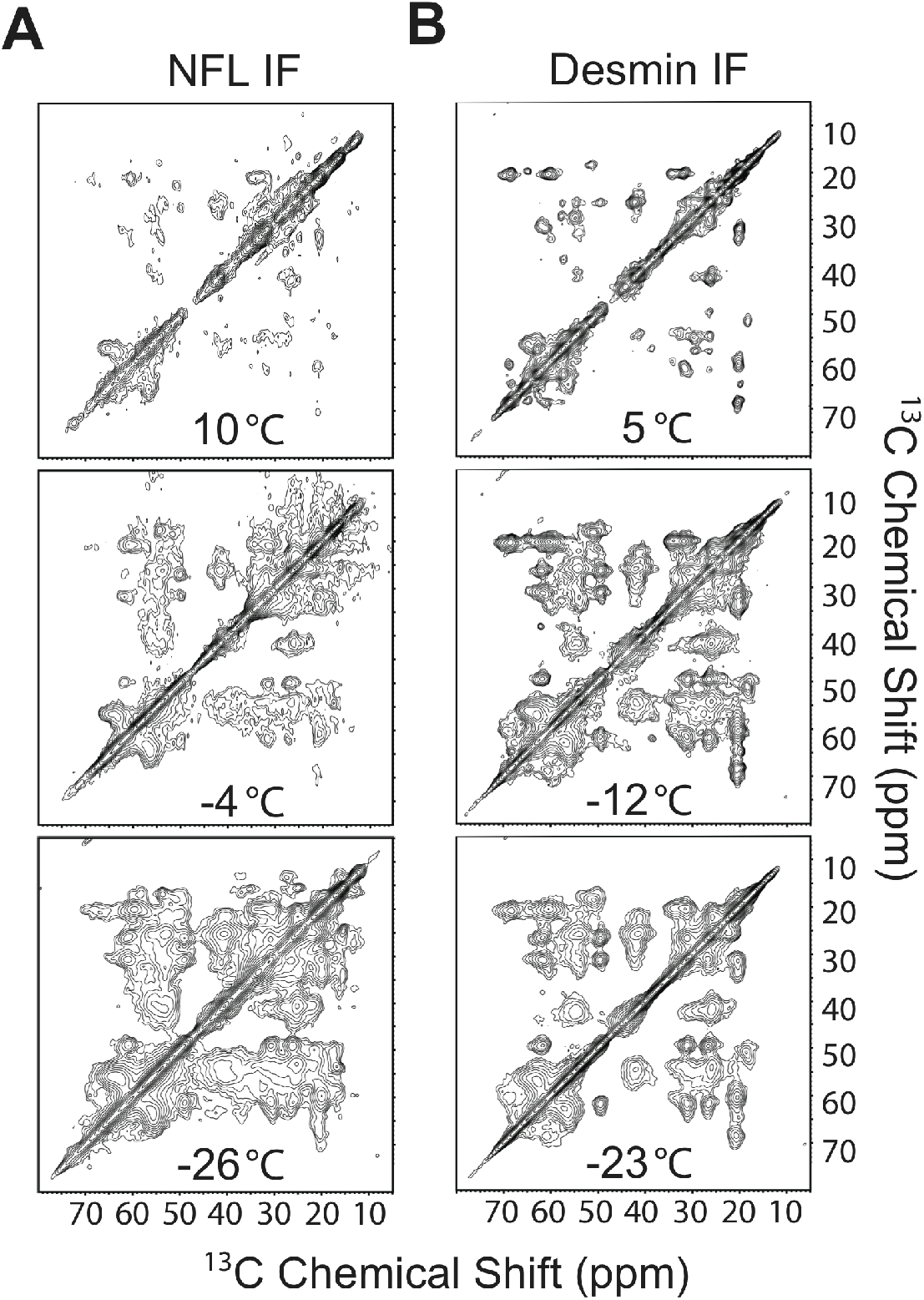
Temperature dependence of two dimensional ^13^C-^13^C ss-NMR spectra of segmentally-labeled intermediate filaments. **(A)** 2D ^13^C-^13^C spectra of ^13^C/^15^N segmentally-labeled NFL head domain embedded within assembled intermediate filaments were recorded at indicated temperatures. Plot counter increment is 1.4 for all spectra. **(B)** 2D ^13^C-^13^C spectra of ^13^C/^15^N segmentally-labeled desmin head domain embedded within assembled intermediate filaments were recorded at indicated temperatures. Plot counter increment is 1.4 for all spectra.

**Figure S5.**
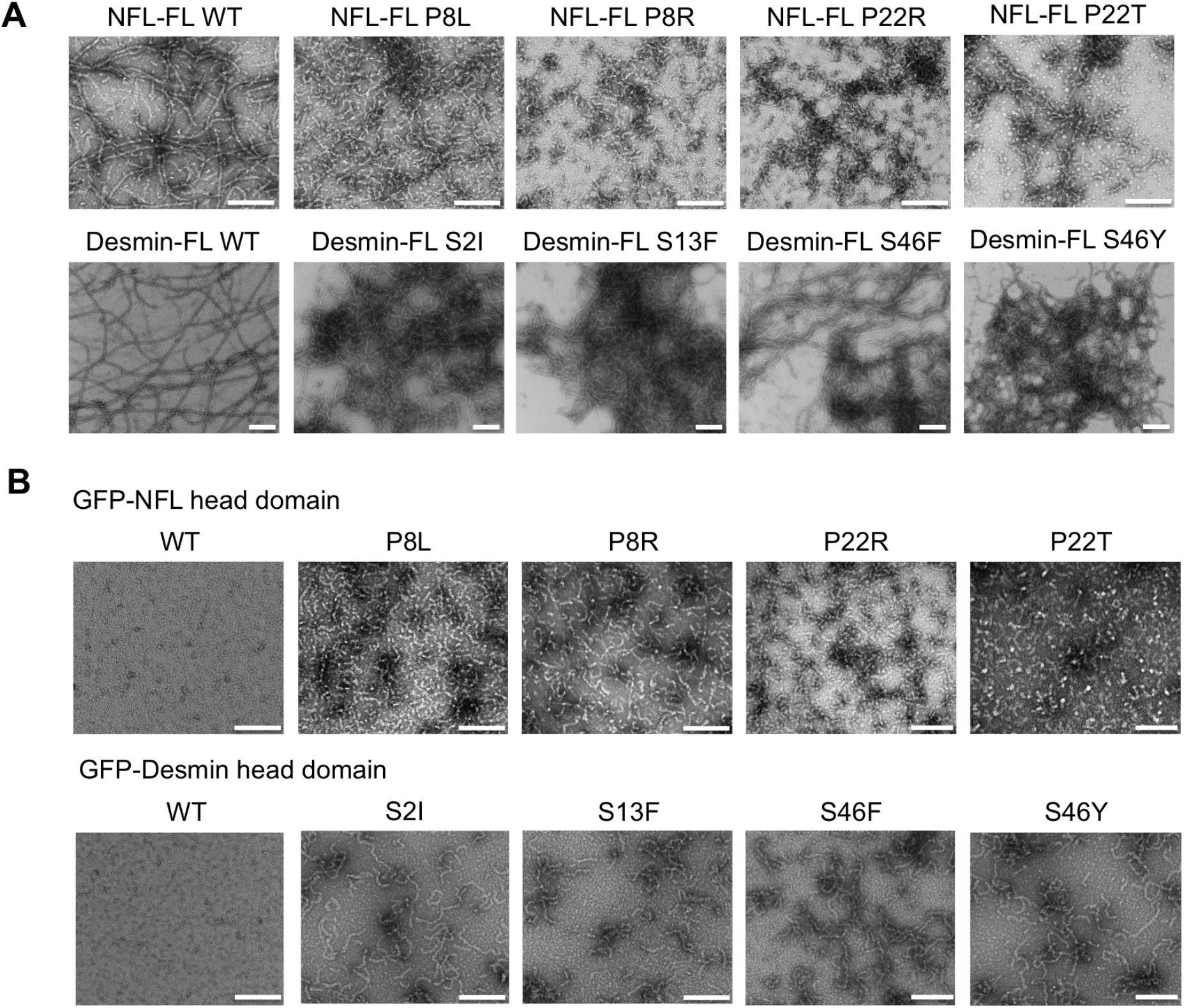
Assays of native and disease-causing mutations in NFL and desmin head domains for assembly into intermediate filaments and head domain-alone polymers. **(A)** NFL full-length (NFL-FL) wild type (WT) protein and indicated head domain mutants were subjected to conditions optimal for intermediate filament assembly (upper panel). All NFL mutants failed to form properly organized intermediate filaments. Desmin full-length (desmin-FL) wild type (WT) and indicated head domain mutants were subjected to conditions optimal for intermediate filament assembly. All desmin mutants formed tangled, aggregated intermediate filaments. (**B**) NFL head domain-only GFP fusions were prepared for the native protein (WT) and indicated head domain mutants. No evidence of polymer formation was observed for the native NFL head domain in 1 M urea. Under the same conditions of incubation, all four NFL head domain mutants formed distinct polymers (upper panel). Desmin head domain-only GFP fusions were prepared for the native protein (WT) and indicated head domain mutants. No evidence of polymer formation was observed for the native desmin head domain in 0.5 M urea. Under the same conditions of incubation, all four desmin head domain mutants formed distinct polymers (lower panel). All scale bars = 200 nm.

**Table S1.**
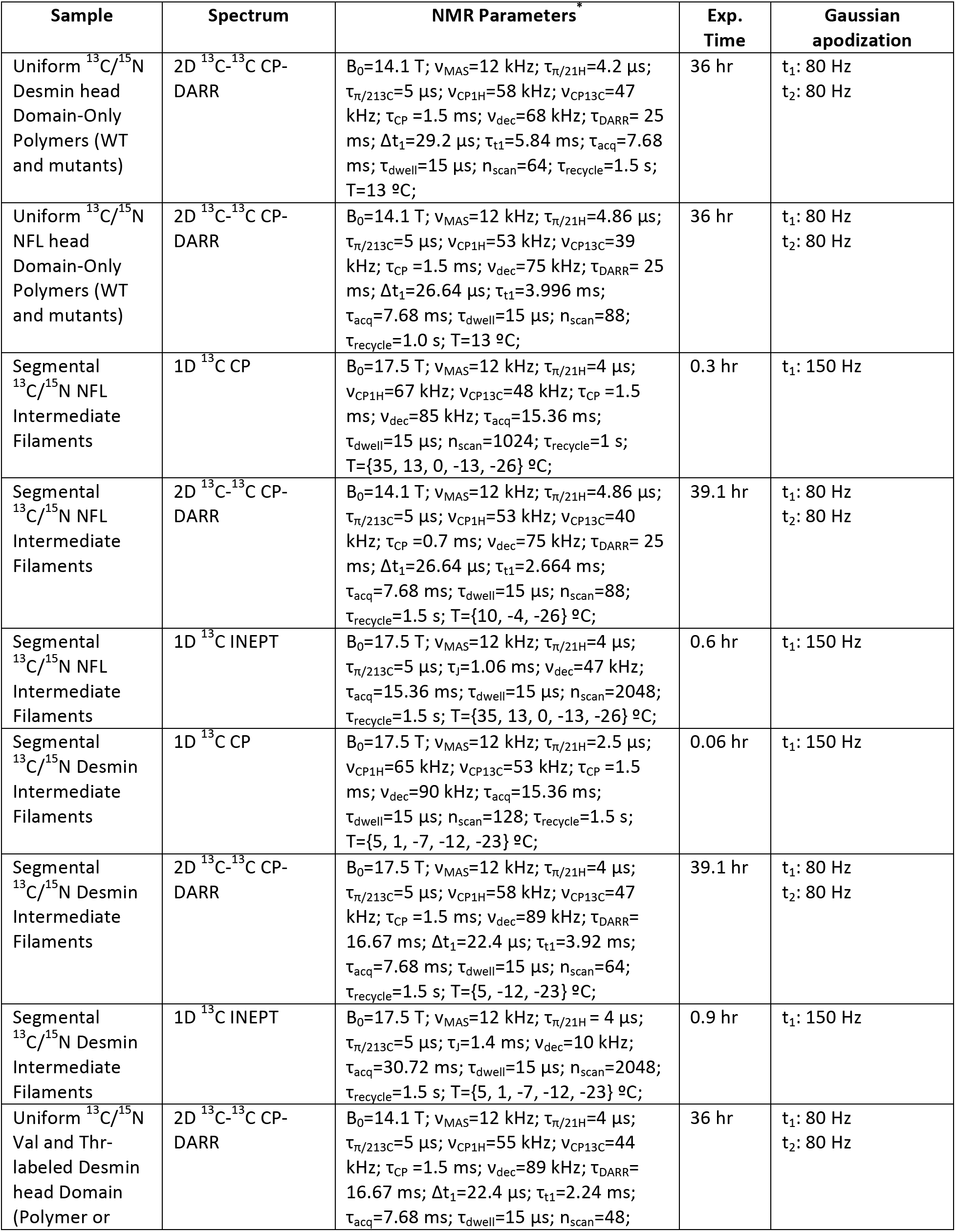
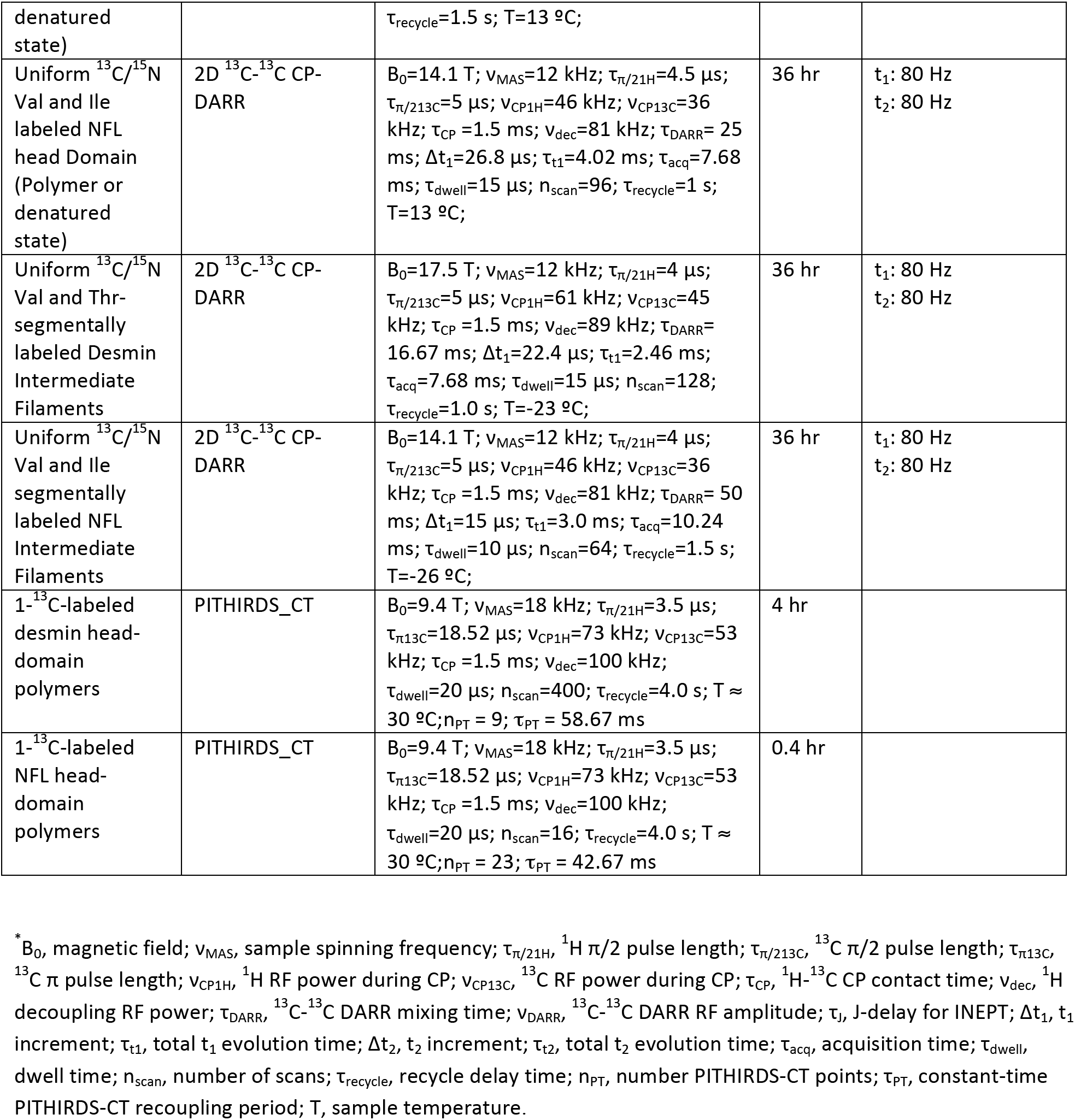
NMR experiments and parameter values for the data presented in this paper.

| Sample | Spectrum | NMR Parameters | Exp. Time | Gaussian apodization |
| --- | --- | --- | --- | --- |
| Uniform $^{13}\text{C}/^{15}\text{N}$ Desmin head Domain-Only Polymers (WT and mutants) | 2D $^{13}\text{C}$-$^{13}\text{C}$ CP-DARR | $B_0=14.1\text{ T}$; $\nu_{\text{MAS}}=12\text{ kHz}$; $\tau_{\pi/21\text{H}}=4.2\text{ }\mu\text{s}$; $\tau_{\pi/213\text{C}}=5\text{ }\mu\text{s}$; $\nu_{\text{CP1H}}=58\text{ kHz}$; $\nu_{\text{CP13C}}=47\text{ kHz}$; $\tau_{\text{CP}}=1.5\text{ ms}$; $\nu_{\text{dec}}=68\text{ kHz}$; $\tau_{\text{DARR}}=25\text{ ms}$; $\Delta t_1=29.2\text{ }\mu\text{s}$; $\tau_{t1}=5.84\text{ ms}$; $\tau_{\text{acq}}=7.68\text{ ms}$; $\tau_{\text{dwell}}=15\text{ }\mu\text{s}$; $n_{\text{scan}}=64$; $\tau_{\text{recycle}}=1.5\text{ s}$; $T=13\text{ }^\circ\text{C}$; | 36 hr | $t_1: 80\text{ Hz}$<br>$t_2: 80\text{ Hz}$ |
| Uniform $^{13}\text{C}/^{15}\text{N}$ NFL head Domain-Only Polymers (WT and mutants) | 2D $^{13}\text{C}$-$^{13}\text{C}$ CP-DARR | $B_0=14.1\text{ T}$; $\nu_{\text{MAS}}=12\text{ kHz}$; $\tau_{\pi/21\text{H}}=4.86\text{ }\mu\text{s}$; $\tau_{\pi/213\text{C}}=5\text{ }\mu\text{s}$; $\nu_{\text{CP1H}}=53\text{ kHz}$; $\nu_{\text{CP13C}}=39\text{ kHz}$; $\tau_{\text{CP}}=1.5\text{ ms}$; $\nu_{\text{dec}}=75\text{ kHz}$; $\tau_{\text{DARR}}=25\text{ ms}$; $\Delta t_1=26.64\text{ }\mu\text{s}$; $\tau_{t1}=3.996\text{ ms}$; $\tau_{\text{acq}}=7.68\text{ ms}$; $\tau_{\text{dwell}}=15\text{ }\mu\text{s}$; $n_{\text{scan}}=88$; $\tau_{\text{recycle}}=1.0\text{ s}$; $T=13\text{ }^\circ\text{C}$; | 36 hr | $t_1: 80\text{ Hz}$<br>$t_2: 80\text{ Hz}$ |
| Segmental $^{13}\text{C}/^{15}\text{N}$ NFL Intermediate Filaments | 1D $^{13}\text{C}$ CP | $B_0=17.5\text{ T}$; $\nu_{\text{MAS}}=12\text{ kHz}$; $\tau_{\pi/21\text{H}}=4\text{ }\mu\text{s}$; $\nu_{\text{CP1H}}=67\text{ kHz}$; $\nu_{\text{CP13C}}=48\text{ kHz}$; $\tau_{\text{CP}}=1.5\text{ ms}$; $\nu_{\text{dec}}=85\text{ kHz}$; $\tau_{\text{acq}}=15.36\text{ ms}$; $\tau_{\text{dwell}}=15\text{ }\mu\text{s}$; $n_{\text{scan}}=1024$; $\tau_{\text{recycle}}=1\text{ s}$; $T=\{35, 13, 0, -13, -26\}\text{ }^\circ\text{C}$; | 0.3 hr | $t_1: 150\text{ Hz}$ |
| Segmental $^{13}\text{C}/^{15}\text{N}$ NFL Intermediate Filaments | 2D $^{13}\text{C}$-$^{13}\text{C}$ CP-DARR | $B_0=14.1\text{ T}$; $\nu_{\text{MAS}}=12\text{ kHz}$; $\tau_{\pi/21\text{H}}=4.86\text{ }\mu\text{s}$; $\tau_{\pi/213\text{C}}=5\text{ }\mu\text{s}$; $\nu_{\text{CP1H}}=53\text{ kHz}$; $\nu_{\text{CP13C}}=40\text{ kHz}$; $\tau_{\text{CP}}=0.7\text{ ms}$; $\nu_{\text{dec}}=75\text{ kHz}$; $\tau_{\text{DARR}}=25\text{ ms}$; $\Delta t_1=26.64\text{ }\mu\text{s}$; $\tau_{t1}=2.664\text{ ms}$; $\tau_{\text{acq}}=7.68\text{ ms}$; $\tau_{\text{dwell}}=15\text{ }\mu\text{s}$; $n_{\text{scan}}=88$; $\tau_{\text{recycle}}=1.5\text{ s}$; $T=\{10, -4, -26\}\text{ }^\circ\text{C}$; | 39.1 hr | $t_1: 80\text{ Hz}$<br>$t_2: 80\text{ Hz}$ |
| Segmental $^{13}\text{C}/^{15}\text{N}$ NFL Intermediate Filaments | 1D $^{13}\text{C}$ INEPT | $B_0=17.5\text{ T}$; $\nu_{\text{MAS}}=12\text{ kHz}$; $\tau_{\pi/21\text{H}}=4\text{ }\mu\text{s}$; $\tau_{\pi/213\text{C}}=5\text{ }\mu\text{s}$; $\tau_j=1.06\text{ ms}$; $\nu_{\text{dec}}=47\text{ kHz}$; $\tau_{\text{acq}}=15.36\text{ ms}$; $\tau_{\text{dwell}}=15\text{ }\mu\text{s}$; $n_{\text{scan}}=2048$; $\tau_{\text{recycle}}=1.5\text{ s}$; $T=\{35, 13, 0, -13, -26\}\text{ }^\circ\text{C}$; | 0.6 hr | $t_1: 150\text{ Hz}$ |
| Segmental $^{13}\text{C}/^{15}\text{N}$ Desmin Intermediate Filaments | 1D $^{13}\text{C}$ CP | $B_0=17.5\text{ T}$; $\nu_{\text{MAS}}=12\text{ kHz}$; $\tau_{\pi/21\text{H}}=2.5\text{ }\mu\text{s}$; $\nu_{\text{CP1H}}=65\text{ kHz}$; $\nu_{\text{CP13C}}=53\text{ kHz}$; $\tau_{\text{CP}}=1.5\text{ ms}$; $\nu_{\text{dec}}=90\text{ kHz}$; $\tau_{\text{acq}}=15.36\text{ ms}$; $\tau_{\text{dwell}}=15\text{ }\mu\text{s}$; $n_{\text{scan}}=128$; $\tau_{\text{recycle}}=1.5\text{ s}$; $T=\{5, 1, -7, -12, -23\}\text{ }^\circ\text{C}$; | 0.06 hr | $t_1: 150\text{ Hz}$ |
| Segmental $^{13}\text{C}/^{15}\text{N}$ Desmin Intermediate Filaments | 2D $^{13}\text{C}$-$^{13}\text{C}$ CP-DARR | $B_0=17.5\text{ T}$; $\nu_{\text{MAS}}=12\text{ kHz}$; $\tau_{\pi/21\text{H}}=4\text{ }\mu\text{s}$; $\tau_{\pi/213\text{C}}=5\text{ }\mu\text{s}$; $\nu_{\text{CP1H}}=58\text{ kHz}$; $\nu_{\text{CP13C}}=47\text{ kHz}$; $\tau_{\text{CP}}=1.5\text{ ms}$; $\nu_{\text{dec}}=89\text{ kHz}$; $\tau_{\text{DARR}}=16.67\text{ ms}$; $\Delta t_1=22.4\text{ }\mu\text{s}$; $\tau_{t1}=3.92\text{ ms}$; $\tau_{\text{acq}}=7.68\text{ ms}$; $\tau_{\text{dwell}}=15\text{ }\mu\text{s}$; $n_{\text{scan}}=64$; $\tau_{\text{recycle}}=1.5\text{ s}$; $T=\{5, -12, -23\}\text{ }^\circ\text{C}$; | 39.1 hr | $t_1: 80\text{ Hz}$<br>$t_2: 80\text{ Hz}$ |
| Segmental $^{13}\text{C}/^{15}\text{N}$ Desmin Intermediate Filaments | 1D $^{13}\text{C}$ INEPT | $B_0=17.5\text{ T}$; $\nu_{\text{MAS}}=12\text{ kHz}$; $\tau_{\pi/21\text{H}}=4\text{ }\mu\text{s}$; $\tau_{\pi/213\text{C}}=5\text{ }\mu\text{s}$; $\tau_j=1.4\text{ ms}$; $\nu_{\text{dec}}=10\text{ kHz}$; $\tau_{\text{acq}}=30.72\text{ ms}$; $\tau_{\text{dwell}}=15\text{ }\mu\text{s}$; $n_{\text{scan}}=2048$; $\tau_{\text{recycle}}=1.5\text{ s}$; $T=\{5, 1, -7, -12, -23\}\text{ }^\circ\text{C}$; | 0.9 hr | $t_1: 150\text{ Hz}$ |
| Uniform $^{13}\text{C}/^{15}\text{N}$ Val and Thr-labeled Desmin head Domain (Polymer or | 2D $^{13}\text{C}$-$^{13}\text{C}$ CP-DARR | $B_0=14.1\text{ T}$; $\nu_{\text{MAS}}=12\text{ kHz}$; $\tau_{\pi/21\text{H}}=4\text{ }\mu\text{s}$; $\tau_{\pi/213\text{C}}=5\text{ }\mu\text{s}$; $\nu_{\text{CP1H}}=55\text{ kHz}$; $\nu_{\text{CP13C}}=44\text{ kHz}$; $\tau_{\text{CP}}=1.5\text{ ms}$; $\nu_{\text{dec}}=89\text{ kHz}$; $\tau_{\text{DARR}}=16.67\text{ ms}$; $\Delta t_1=22.4\text{ }\mu\text{s}$; $\tau_{t1}=2.24\text{ ms}$; $\tau_{\text{acq}}=7.68\text{ ms}$; $\tau_{\text{dwell}}=15\text{ }\mu\text{s}$; $n_{\text{scan}}=48$; | 36 hr | $t_1: 80\text{ Hz}$<br>$t_2: 80\text{ Hz}$ |
| denatured state) | | $\tau_{\text{recycle}}=1.5 \text{ s}$; $T=13 \text{ }^{\circ}\text{C}$; | | |
| Uniform $^{13}\text{C}/^{15}\text{N}$ Val and Ile labeled NFL head Domain (Polymer or denatured state) | 2D $^{13}\text{C}$-$^{13}\text{C}$ CP-DARR | $B_0=14.1 \text{ T}$; $\nu_{\text{MAS}}=12 \text{ kHz}$; $\tau_{\pi/21\text{H}}=4.5 \text{ }\mu\text{s}$; $\tau_{\pi/213\text{C}}=5 \text{ }\mu\text{s}$; $\nu_{\text{CP1H}}=46 \text{ kHz}$; $\nu_{\text{CP13C}}=36 \text{ kHz}$; $\tau_{\text{CP}}=1.5 \text{ ms}$; $\nu_{\text{dec}}=81 \text{ kHz}$; $\tau_{\text{DARR}}=25 \text{ ms}$; $\Delta t_1=26.8 \text{ }\mu\text{s}$; $\tau_{t1}=4.02 \text{ ms}$; $\tau_{\text{acq}}=7.68 \text{ ms}$; $\tau_{\text{dwell}}=15 \text{ }\mu\text{s}$; $n_{\text{scan}}=96$; $\tau_{\text{recycle}}=1 \text{ s}$; $T=13 \text{ }^{\circ}\text{C}$; | 36 hr | $t_1: 80 \text{ Hz}$<br>$t_2: 80 \text{ Hz}$ |
| Uniform $^{13}\text{C}/^{15}\text{N}$ Val and Thr-segmentally labeled Desmin Intermediate Filaments | 2D $^{13}\text{C}$-$^{13}\text{C}$ CP-DARR | $B_0=17.5 \text{ T}$; $\nu_{\text{MAS}}=12 \text{ kHz}$; $\tau_{\pi/21\text{H}}=4 \text{ }\mu\text{s}$; $\tau_{\pi/213\text{C}}=5 \text{ }\mu\text{s}$; $\nu_{\text{CP1H}}=61 \text{ kHz}$; $\nu_{\text{CP13C}}=45 \text{ kHz}$; $\tau_{\text{CP}}=1.5 \text{ ms}$; $\nu_{\text{dec}}=89 \text{ kHz}$; $\tau_{\text{DARR}}=16.67 \text{ ms}$; $\Delta t_1=22.4 \text{ }\mu\text{s}$; $\tau_{t1}=2.46 \text{ ms}$; $\tau_{\text{acq}}=7.68 \text{ ms}$; $\tau_{\text{dwell}}=15 \text{ }\mu\text{s}$; $n_{\text{scan}}=128$; $\tau_{\text{recycle}}=1.0 \text{ s}$; $T=-23 \text{ }^{\circ}\text{C}$; | 36 hr | $t_1: 80 \text{ Hz}$<br>$t_2: 80 \text{ Hz}$ |
| Uniform $^{13}\text{C}/^{15}\text{N}$ Val and Ile segmentally labeled NFL Intermediate Filaments | 2D $^{13}\text{C}$-$^{13}\text{C}$ CP-DARR | $B_0=14.1 \text{ T}$; $\nu_{\text{MAS}}=12 \text{ kHz}$; $\tau_{\pi/21\text{H}}=4 \text{ }\mu\text{s}$; $\tau_{\pi/213\text{C}}=5 \text{ }\mu\text{s}$; $\nu_{\text{CP1H}}=46 \text{ kHz}$; $\nu_{\text{CP13C}}=36 \text{ kHz}$; $\tau_{\text{CP}}=1.5 \text{ ms}$; $\nu_{\text{dec}}=81 \text{ kHz}$; $\tau_{\text{DARR}}=50 \text{ ms}$; $\Delta t_1=15 \text{ }\mu\text{s}$; $\tau_{t1}=3.0 \text{ ms}$; $\tau_{\text{acq}}=10.24 \text{ ms}$; $\tau_{\text{dwell}}=10 \text{ }\mu\text{s}$; $n_{\text{scan}}=64$; $\tau_{\text{recycle}}=1.5 \text{ s}$; $T=-26 \text{ }^{\circ}\text{C}$; | 36 hr | $t_1: 80 \text{ Hz}$<br>$t_2: 80 \text{ Hz}$ |
| 1-$^{13}\text{C}$-labeled desmin head-domain polymers | PITHIRDS_CT | $B_0=9.4 \text{ T}$; $\nu_{\text{MAS}}=18 \text{ kHz}$; $\tau_{\pi/21\text{H}}=3.5 \text{ }\mu\text{s}$; $\tau_{\pi13\text{C}}=18.52 \text{ }\mu\text{s}$; $\nu_{\text{CP1H}}=73 \text{ kHz}$; $\nu_{\text{CP13C}}=53 \text{ kHz}$; $\tau_{\text{CP}}=1.5 \text{ ms}$; $\nu_{\text{dec}}=100 \text{ kHz}$; $\tau_{\text{dwell}}=20 \text{ }\mu\text{s}$; $n_{\text{scan}}=400$; $\tau_{\text{recycle}}=4.0 \text{ s}$; $T \approx 30 \text{ }^{\circ}\text{C}$; $n_{\text{PT}}=9$; $\tau_{\text{PT}}=58.67 \text{ ms}$ | 4 hr | |
| 1-$^{13}\text{C}$-labeled NFL head-domain polymers | PITHIRDS_CT | $B_0=9.4 \text{ T}$; $\nu_{\text{MAS}}=18 \text{ kHz}$; $\tau_{\pi/21\text{H}}=3.5 \text{ }\mu\text{s}$; $\tau_{\pi13\text{C}}=18.52 \text{ }\mu\text{s}$; $\nu_{\text{CP1H}}=73 \text{ kHz}$; $\nu_{\text{CP13C}}=53 \text{ kHz}$; $\tau_{\text{CP}}=1.5 \text{ ms}$; $\nu_{\text{dec}}=100 \text{ kHz}$; $\tau_{\text{dwell}}=20 \text{ }\mu\text{s}$; $n_{\text{scan}}=16$; $\tau_{\text{recycle}}=4.0 \text{ s}$; $T \approx 30 \text{ }^{\circ}\text{C}$; $n_{\text{PT}}=23$; $\tau_{\text{PT}}=42.67 \text{ ms}$ | 0.4 hr | |

\* $B_0$ , magnetic field; $\nu_{\text{MAS}}$ , sample spinning frequency; $\tau_{\pi/21\text{H}}$ , $^1\text{H}$ $\pi/2$ pulse length; $\tau_{\pi/213\text{C}}$ , $^{13}\text{C}$ $\pi/2$ pulse length; $\tau_{\pi13\text{C}}$ , $^{13}\text{C}$ $\pi$ pulse length; $\nu_{\text{CP1H}}$ , $^1\text{H}$ RF power during CP; $\nu_{\text{CP13C}}$ , $^{13}\text{C}$ RF power during CP; $\tau_{\text{CP}}$ , $^1\text{H}$ - $^{13}\text{C}$ CP contact time; $\nu_{\text{dec}}$ , $^1\text{H}$ decoupling RF power; $\tau_{\text{DARR}}$ , $^{13}\text{C}$ - $^{13}\text{C}$ DARR mixing time; $\nu_{\text{DARR}}$ , $^{13}\text{C}$ - $^{13}\text{C}$ DARR RF amplitude; $\tau_{\text{J}}$ , J-delay for INEPT; $\Delta t_1$ , $t_1$ increment; $\tau_{t1}$ , total $t_1$ evolution time; $\Delta t_2$ , $t_2$ increment; $\tau_{t2}$ , total $t_2$ evolution time; $\tau_{\text{acq}}$ , acquisition time; $\tau_{\text{dwell}}$ , dwell time; $n_{\text{scan}}$ , number of scans; $\tau_{\text{recycle}}$ , recycle delay time; $n_{\text{PT}}$ , number PITHIRDS-CT points; $\tau_{\text{PT}}$ , constant-time PITHIRDS-CT recoupling period; $T$ , sample temperature.

## Materials and methods

### Protein expression and purification

Standard molecular cloning strategies were employed to generate expression plasmids for all recombinant protein constructs. Briefly, neurofilament light (NFL) and desmin genes were cloned from a human cDNA library. Sequences corresponding to full-length proteins and truncated variants were then amplified by PCR and cloned into pHis-parallel vectors to produce recombinant constructs harboring N-terminal fusion tags (6xHis, 6xHis-GFP, and 6xHis-mCherry). To enable segmental isotopic labeling via protein splicing, chimeras consisting of NFL or desmin constructs fused to the Cfa split intein or intact Cfa intein (Stevens et al., 2016) were assembled via the Gibson Assembly protocol (detailed construct information described below). All mutation constructs were produced via standard site-directed mutagenesis protocols. All cloning and mutagenesis was confirmed by Sanger sequencing (Eurofins Genomics).

Recombinant proteins were expressed in *E. coli* BL21 (DE3) cells grown to an OD_600_ of 0.6 in LB medium. Expression of full-length NFL, full-length desmin, 6xHis-tagged NFL head domain, and 6xHis-tagged desmin head domain was induced via addition of 0.8 mM IPTG at 37 °C and cells were harvested 4 hours post-induction. Expression of all GFP and mCherry fusion proteins was induced with 0.6 mM IPTG at 16 °C and cells were harvested 16 hours post-induction. Universal and amino acid-specific ^13^C, ^15^N labeling of NFL and desmin head domains was carried out as previously described (Murray et al., 2017; Murray et al., 2018).

For purification of full-length NFL, full-length desmin, 6xHis-tagged NFL head domain, and 6xHis-tagged desmin head domain, cell pellets were resuspended in a lysis buffer containing 25 mM Tris-HCl (pH 7.5), 200 mM NaCl, 10 mM β-mercaptoethanol (β-ME), and protease inhibitors (EDTA-free protease inhibitor cocktail tablet, Sigma) and disrupted via sonication for 3 minutes (Fisher Scientific, 10 seconds on/30 seconds off, 78% power). 6xHis-tagged NFL head domain and 6xHis-tagged desmin head domain inclusion bodies were collected by centrifugation at 3,000 x g for 30 minutes. The inclusion body pellet was resuspended in solubilization buffer containing 25 mM Tris-HCl (pH 7.5), 200 mM NaCl, 8 M Urea, and 10 mM β-ME, sonicated for 1 minute (10 seconds on/30 seconds off, 78% power), and clarified via centrifugation at 36,000 x g for 50 minutes. Supernatant was then applied to Ni^2+^-NTA resin (Qiagen) and the column washed with a solubilization buffer supplemented with 20 mM imidazole and eluted in solubilization buffer supplemented with 300 mM imidazole. Pure protein was concentrated by centrifugal filtration (Amicon Ultra-15) and samples were aliquoted, flash frozen, and stored at −80 °C for future use.

Full-length NFL and desmin inclusion bodies were collected by centrifugation at 36,000 x g for 20 minutes. The inclusion body pellet was resuspended in solubilization buffer lacking NaCl and sonicated for 1 minute (10 seconds on/30 seconds off, 78% power). This solution was then clarified via centrifugation at 36,000 x g for 50 minutes and the supernatant was passed through 0.22 μm filter (Millex GV) and loaded onto Hitrap Q column (GE Healthcare). A salt gradient elution was then performed (0 – 0.5 M NaCl in solubilization buffer) and elution fractions were analyzed by SDS-PAGE. >Pure protein was concentrated by centrifugal filtration (Amicon Ultra-15) and samples were aliquoted, flash frozen, and stored at −80 °C for future use.

For purification of GFP- and mCherry-fused NFL and desmin proteins, cells were resuspended in a lysis buffer containing 25 mM Tris-HCl (pH 7.5), 200 mM NaCl, 2 M urea, 10 mM β-ME, 20 mM imidazole, and protease inhibitors and disrupted by sonication for 3 minutes (10 seconds on/30 seconds off, 78% power). Cell lysates were clarified via centrifugation at 36,000 x g for 50 minutes and supernatant was applied to Ni^2+^-NTA resin, washed with lysis buffer, and bound proteins were eluted in lysis buffer supplemented with 300 mM imidazole. Eluted protein was then diluted 5-fold with low salt ion-exchange buffer (25 mM Tris-HCl (pH 7.5), 100 mM NaCl, 2 M urea, and 2 mM tris(2-carboxyethyl)phosphine (TCEP)) and applied to a HiTrap Q column. A salt gradient elution was then performed (0 – 0.5 M NaCl in ion-exchange buffer) and elution fractions were analyzed by SDS-PAGE. Pure protein was concentrated by centrifugal filtration (Amicon Ultra-15) and samples were aliquoted, flash frozen, and stored at −80 °C for future use.

### Preparation of NFL and desmin polymers and hydrogels

6xHis-tagged NFL head domain, 6xHis-tagged desmin head domain and their respective mutants were diluted to 100 - 200 μM in gelation buffer (25 mM Tris-HCl (pH 7.5), 200 mM NaCl, and 10 mM β-ME) supplemented with 8 M urea and dialyzed stepwise at 25 °C against gelation buffer containing 4 M urea, 2 M urea, 1 M urea, 0.5 M urea, and no urea to remove denaturant and initiate polymer formation.

Preparation of GFP-fused NFL head domain hydrogel droplets was achieved by dialyzing the purified GFP-NFL head domain against a gelation buffer supplemented with 0.5 mM EDTA, and protease inhibitors for 16 hours at 25 °C. Dialyzed protein was concentrated to roughly 20 mg/mL and sonicated for 3 seconds at 1% power. Following centrifugation, 20 μL droplets were deposited onto a piece of parafilm fixed inside a 10 cm cell culture dish. The dish was sealed with parafilm and incubated for 16 hours at 25 °C.

The mCherry-fused NFL head domain and desmin head domain were diluted to 200 μM and dialyzed against gelation buffer for 16 hours at 25 °C. Precipitates were removed by centrifugation at 10,000 x g for 1 minute. Head domain polymers, which remain in the supernatant, were analyzed by negative staining transmission electron microscopy (TEM). Hydrogel droplets were prepared as previously described (Kato et al., 2017). Briefly, 0.5 μL of the solution containing mCherry-fused head domain polymers was deposited onto a 35 mm glass bottom dish containing a piece of filter paper soaked with 35 μL of the gelation buffer. The dish was sealed with parafilm and incubated overnight at 25 °C.

To compare polymer formation propensity between wild-type head domains and their mutant variants, the 6xHis-tagged NFL head domain and disease mutants were diluted to 100 μM and dialyzed against gelation buffer supplemented 4 M urea for 16 hours at 25 °C. Similarly, the 6xHis-tagged desmin head domain and disease mutants were diluted to 100 μM and dialyzed against gelation buffer supplemented 3 M urea for 3 hours at 25 °C. For the GFP-tagged NFL head domain and its mutant variants, proteins were diluted to 50 μM and dialyzed against gelation buffer supplemented 1 M urea for 2 hours at 25 °C.

Similarly, GFP-tagged desmin head domain and its mutant variants were diluted to 50 μM and dialyzed against gelation buffer supplemented 0.5 M urea for 16 hours at 25 °C. All head domain polymers were analyzed by negative staining TEM.

### Hydrogel-binding assays

Intact mouse brain tissue was flash frozen in liquid nitrogen and subjected to cryogenic grinding (cryomill, Retsch). Resultant frozen powder was resuspended at 100 mg/mL in gelation buffer and clarified via centrifugation at 12,000 x g for 10 minutes at 4 °C. 1.2 mL of supernatant (mouse brain lysate) was then incubated with ten GFP-NFL head domain hydrogel droplets for 2 hours at 4 °C. Following incubation, the droplets were spun down at 800 x g for 5 minutes at 4 °C, washed three times with gelation buffer supplemented with 0.5 mM EDTA and protease inhibitors, and melted in gelation buffer supplemented with 6 M guanidine-HCl. Solubilized protein solution was then applied to Ni^2+^-NTA resin to fully capture and remove the bait protein (6xHis-GFP-NFL head domain). Flow-through consisting of hydrogel-interacting proteins was collected and analyzed via mass spectrometry-based shotgun proteomics.

To map the domains of NFL and desmin that mediate self-association with their respective head domain hydrogel droplets, the GFP-labeled variants described in Figure 2 were purified as described above. GFP-labeled proteins were diluted to 1 μM with 1 mL of gelation buffer immediately added to the corresponding mCherry-fused head domain hydrogel droplets for 12 hours. After incubation, the hydrogel droplets were washed twice with gelation buffer to remove unbound protein prior to confocal microscopy imaging analysis (Kato et al., 2017).

### Intermediate filament assembly

To assemble NFL intermediate filaments (IFs), full-length wild-type NFL or mutant variants thereof were diluted to 10 μM with a buffer containing 25 mM Tris-HCl (pH 7.5), 200 mM NaCl, 8 M urea, 1 mM TCEP, 0.5 mM EDTA and dialyzed against a buffer containing 50 mM MES (pH 6.25), 170 mM NaCl, 1 mM TCEP, and 1 mM EGTA for 16 hours at 25 °C.

To assemble desmin IFs, full-length wild-type desmin or mutant variants thereof were diluted to 20 μM with a buffer containing 25 mM Tris-HCl (pH 7.5), 200 mM NaCl, 8 M urea, 1 mM TCEP, and 0.5 mM EDTA and dialyzed stepwise against a buffer containing 5 mM Tris-HCl (pH 8.5), 1 mM EDTA, 0.1 mM EGTA, and 10 mM β-ME and supplemented with 4 M urea, 2 M urea, and no urea to remove denaturant. Dialyzed desmin proteins were then mixed with a buffer containing 45 mM Tris-HCl (pH 7.0) and 100 mM NaCl at a 1:1 ratio by volume and incubated for 1 hour at 25 °C.

### Negative staining transmission electron microscopy

Head domain polymer solution (5 μL) was added to an EM grid for 10 seconds and excess sample was removed via blotting with filter paper. The grid was then washed with water for 2 seconds and stained with 5 μL uranyl acetate (2%) for 15 seconds. An identical protocol was employed to generate EM grids containing assembled IFs except the water washing step was omitted. All negative staining TEM samples were imaged on a JOEL 1400 microscope.

### Intein-mediated segmental isotopic labeling of NFL and desmin proteins

For segmental labeling of NFL, a fusion construct comprising the GFP-NFL head domain (residues 2-85) and a C-terminal CfaN split intein fragment (Stevens et al., 2016) was expressed in minimal media supplemented with the desired ^13^C, ^15^N-labeled amino acids. A complementary fusion construct comprising headless NFL (residues 86-543, I86C) and an N-terminal GFP-CfaC split intein fragment was produced in LB medium as described above. Note that Cfa intein-mediated protein splicing requires a cysteine (I86C) at the ligation junction (Figure S2). Purification of the fusion constructs was carried out as described above. Isotopically-labeled head domain and unlabeled headless domain fusion proteins were then mixed at a 2:1 molar ratio (100 μM: 50 μM) in the presence of 0.05 mg/mL caspase-3 enzyme to remove the GFP tag from each construct. The mixture was then dialyzed against a buffer containing 40 mM KH_2_PO_4_ (pH 7.2), 150 mM NaCl, and 1 mM TCEP at 25 °C for 16 hours and diluted 10-fold into a buffer containing 25 mM Tris-HCl (pH 7.4), 8 M urea, and 1 mM TCEP to solubilize any precipitate that formed during dialysis. Full-length, segmentally-labeled NFL protein was then purified following the HiTrap Q chromatography protocol described above.

For segmental labeling of desmin, a fusion construct comprising the GFP-desmin head domain (residues 2-84) and the contiguous Cfa intein was expressed in minimal media supplemented with the desired ^13^C, ^15^N-labeled amino acids. A complementary unlabeled 6xHis-headless desmin construct (residues 85-470, A85C) was purified as described above. The 6xHis tag was removed upon treatment with the caspase-3 enzyme, liberating a headless desmin construct that harbors a native chemical ligation-compatible N-terminal cysteine (Figure S2). Isotopically-labeled head domain and unlabeled headless domain fusion proteins were then mixed at a 1:1 molar ratio (200 μM: 200 μM) in a buffer containing 100 mM KH_2_PO_4_ (pH 7.2), 200 mM MESNa, 100 mM NaCl, 5 mM TCEP, 1 mM EDTA, 0.05 mg/mL caspase-3 enzyme, and 200 mM sodium 2-mercaptoethanesulfonate (MESNa) for 16 hours at 25 °C. Following incubation, the sample was diluted 10-fold into a buffer containing 25 mM Tris-HCl (pH 7.4), 8 M urea, and 1 mM TCEP. Full-length, segmentally-labeled desmin protein was then purified following the HiTrap Q chromatography protocol described above.

Intermediate filament assembly using the full-length, segmentally-labeled proteins was carried out as described above and subsequent filament compaction was achieved by ultracentrifugation at 100,000 x g for 30 minutes.

### Sample packing for solid state NMR (ss-NMR) measurement

NFL and desmin head-domain polymers and IFs were centrifuged at 247,000 x g for 1 hour and supernatant was discarded. Pellets were then transferred to 0.7 mL ultracentrifuge tubes (Beckman Coulter, part #: 343776), from which they were packed into 3.2 mm-diameter magic-angle spinning (MAS) rotors (22 μL or 36 μL sample volume, Revolution NMR) by centrifugation (swinging bucket) at 50,000 x g for 30 minutes using a home-made device that holds the MAS rotor and the 0.7 mL tube.

After centrifugation, excess buffer was blotted from the MAS rotor and the rotor caps were sealed with cyanoacrylate glue. For the ether precipitated NFL and desmin head domain samples, the head domains in 6 M guanidine-HCl buffer were dialyzed overnight against 2 L of ultrapure water. Resulting precipitates were flash frozen in liquid nitrogen and lyophilized for 3 hours. Lyophilized proteins were then dissolved in 100 μl of trifluoroacetic acid. 1 mL of ice cold tert-butyl methyl ether was then added to the solution, vortexed for 30 seconds, then centrifuged at 21,000 x g for 30 minutes in a swinging bucket rotor pre-cooled to 4 °C. Supernatant was removed and the pellets were dried under a stream of nitrogen gas. Dried pellets were then packed manually into MAS rotors.

PITHIRDS-CT measurements were performed on NFL or desmin head polymer samples in which backbone carbonyl sites of either all Val residues or all Leu residues (in NFL) or either all Val residues, all Leu residues, or all Phe residues (in desmin) were ^13^C-labeled. Samples for these measurements were packed into 3.2 mm MAS rotors as fully hydrated pellets (see above) then lyophilized in the rotors. Lyophilization was performed because the transverse spin relaxation times (T_2_ values) of carbonyl ^13^C labels in the fully hydrated samples were found to be too short (<20 ms) to permit PITHIRDS-CT measurements. Such measurements require constant-time dipolar recoupling periods greater than 30 ms for ^13^C-^13^C distances greater than 5 Å. T_2_ values of lyophilized samples were found to be ~60 ms under the PITHIRDS-CT pulse sequence.

### ss-NMR measurements

1D and 2D ss-NMR spectra were acquired at 17.5 T (746.1 MHz ^1^H NMR frequency) with a Varian Infinity spectrometer console and a 3.2 mm low-E MAS probe produced by Black Fox, Inc. (Tallahassee, FL) or at 14.1 T (599.2 MHz ^1^H NMR frequency) with a Varian InfinityPlus spectrometer console and a 3.2 mm Varian BioMAS probe. PITHIRDS-CT data were acquired at 9.4 T (400.8 MHz ^1^H NMR frequency) with a Bruker Avance III spectrometer console and a 3.2 mm Varian T3 MAS NMR probe. Sample temperatures were controlled by cooled nitrogen gas and determined from the temperature-dependent ^1^H NMR frequency of water within the NFL or desmin sample or the temperature-dependent ^89^Br NMR frequency of an external sample of KBr powder (Thurber and Tycko, 2009). ^13^C-^13^C polarization transfers in 2D spectra used Dipolar-Assisted Rotational Resonance (DARR) (Takegoshi et al., 2001). ^1^H decoupling in CP-based measurements used two-pulse phase modulation (TPPM) (Bennett et al., 1995). ^1^H decoupling in INEPT-based measurements used 25-pulse composite π sequences (Tycko et al., 1985). PITHIRDS-CT measurements used the constant-time homonuclear dipolar recoupling pulse sequence described previously (Tycko, 2007), with pulsed spin-lock detection (Petkova and Tycko, 2002) to enhance sensitivity. Other conditions and parameters for ss-NMR measurements are given in Table S1.

1D NMR data were processed with Varian Spinsight or Bruker TopSpin software. 2D data were processed with NMRPipe software (Delaglio et al., 1995). Pure Gaussian apodization functions were used to process all data, with no artificial resolution enhancement to reduce apparent NMR linewidths. PITHIRDS-CT data were analyzed by comparison with numerical simulations.

### NFL and desmin head domain phosphorylation

As described above GFP-fused NFL head domain and GFP-fused desmin head domain monomers were diluted to 1 μM in gelation buffer and co-incubated with respective mCherry-fused head domain hydrogel droplets for 6 hours at 25 °C. Unbound monomers were removed from hydrogel droplets via gelation buffer washes and droplets were equilibrated in phosphorylation reaction buffer (40 mM Tris-HCl (pH 7.4), 20 mM MgCl_2_). ATP (0.2 mM) and/or protein kinase A (0.1 mg/mL; PKA, Promega) were then added to hydrogel droplets for 12 hours at 25 °C. Post-incubation hydrogel droplets were visualized by confocal microscopy.

Phosphorylation of full-length NFL and desmin IFs was carried out by adding 20 mM MgCl_2_ to the assembled IFs (10 μM) followed by addition of ATP (0.2 mM) and/or PKA (0.1 mg/mL) for 5 hours at 30 °C. The effect of PKA activity on IF structure was analyzed by negative staining TEM.

